# Asymmetric effects of acute stress on cost and benefit learning

**DOI:** 10.1101/2021.04.25.441347

**Authors:** Stella Voulgaropoulou, Fasya Fauzani, Janine Pfirrmann, Claudia Vingerhoets, Thérèse van Amelsvoort, Dennis Hernaus

## Abstract

Stressful events trigger a complex physiological reaction – the *fight-or-flight* response – that can hamper flexible decision-making. Inspired by key neural and peripheral characteristics of the fight-or-flight response, here we ask whether acute stress changes how humans learn about costs and benefits. Participants were randomly exposed to an acute stress or no-stress control condition after which they completed a cost-benefit reinforcement learning task. Acute stress improved learning to maximize benefits (monetary rewards) relative to minimising energy expenditure (grip force). Using computational modelling, we demonstrate that costs and benefits can exert asymmetric effects on decisions when prediction errors that convey information about the reward value and cost of actions receive inappropriate importance; a process associated with distinct alterations in pupil size fluctuations. These results provide new insights into learning strategies under acute stress – which, depending on the context, may be maladaptive or beneficial - and candidate neuromodulatory mechanisms that could underlie such behaviour.

## Introduction

Stress is ubiquitous in everyday life. From recurrent, brief, events (a work meeting, moving to a new house) to major life events (armed combat, pandemic, financial crisis), humans are continuously exposed to challenges in their daily environment. The immediate central and peripheral physiological cascade triggered by such events, collectively termed the *fight-or-flight (or acute stress) response* (Cannon, 1915), serves an allostatic role that enables organisms to adequately respond to environmental demands (de Kloet, Joëls, & Holsboer, 2005). Although beneficial for survival, this allostatic process comes at a cost: stress-induced redistributions of neural resources - e.g., towards vigilance or threat detection - may hamper the deployment of strategies that support adaptive and optimal decision-making (Hermans, Henckens, Joëls, & Fernández, 2014).

Optimal decisions essentially depend on the ability to rapidly learn from the positive and negative outcomes of previous actions, also known as *reinforcement learning* (Niv, 2009). Considerable evidence now suggests that acute stress impairs aspects of reinforcement learning (Carvalheiro, Conceição, Mesquita, & Seara-Cardoso, 2020; de Berker et al., 2016; Raio, Hartley, Orederu, Li, & Phelps, 2017). Acute stress, among others, modulates the impact of positive outcomes on future decisions - both positively and negatively - (Berghorst, Bogdan, Frank, & Pizzagalli, 2013; Carvalheiro et al., 2020; Lighthall, Gorlick, Schoeke, Frank, & Mather, 2013; Petzold, Plessow, Goschke, & Kirschbaum, 2010), likely driven by changes in reward sensitivity and the signalling of reward prediction errors (RPEs) (Berghorst et al., 2013; Carvalheiro et al., 2020; Huys, Pizzagalli, Bogdan, & Dayan, 2013); putatively dopaminergic teaching signals that represent the mismatch between actual and expected outcomes, which are used to flexibly adjust behaviour (Niv, 2009; Rescorla, 1972). Alterations in the influence of RPEs on future decisions play a key role in the development of motivational impairments, which are frequently observed in behavioural disorders associated with repeated and/or prolonged stress exposure (Huys et al., 2013).

Intuitive as it is, the notion that the impact of acute stress on (potentially maladaptive) decisions *primarily* involves changes in how reward value influences action may be oversimplified. Decisions are not only motivated by appetitive properties; they equally depend on the – cognitive (e.g., mental effort) or physical (e.g., energy) – *cost* associated with actions (Hauser, Eldar, & Dolan, 2017; Pessiglione, Vinckier, Bouret, Daunizeau, & Le Bouc, 2017; Schmidt, Lebreton, Cléry-Melin, Daunizeau, & Pessiglione, 2012). Expectations about action costs are also updated according to a prediction error rule (Skvortsova, Degos, Welter, Vidailhet, & Pessiglione, 2017; Skvortsova, Palminteri, & Pessiglione, 2014) (henceforth “effort” prediction errors; EPEs), which due to the aversive and resource-consuming nature of effort, optimal learners should utilize to minimize effort expenditure. When decisions involve a potential cost *and* benefit, the former is subtracted from the latter to compute a “net” or subjective decision value (i.e., effort-discounted reward value) (Klein-Flügge, Kennerley, Friston, & Bestmann, 2016; Skvortsova et al., 2017; Skvortsova et al., 2014). Notably, stress exposure impairs cost-benefit decisions in rodents when learning is not explicitly required (Friedman et al., 2017; Shafiei, Gray, Viau, & Floresco, 2012). Moreover, in a reinforcement learning context, acute stress blocks the flexible updating of aversive value (Raio et al., 2017), an inherent property of costly actions. These results suggest that decisions during acute stress may involve a complex shift in reinforcement learning strategies that serve to balance the cost versus benefits of decisions; a hypothesis that hitherto has remained unexplored.

Although computationally similar in nature, distinct neural correlates of RPEs (e.g., striatal subdivisions, ventromedial prefrontal cortex [vmPFC]) and EPEs (e.g., parietal cortex, insula, dorsomedial PFC) can be observed in cost-benefit reinforcement learning paradigms (Hauser et al., 2017; Skvortsova et al., 2014). The ascending dopaminergic (e.g., RPEs, action cost, reward value) (Schultz, Dayan, & Montague, 1997; Skvortsova et al., 2017; Yohn et al., 2016), noradrenergic (e.g., mobilizing energy) (Pessiglione et al., 2017; Varazzani, San-Galli, Gilardeau, & Bouret, 2015) and serotonergic (e.g., aversive value, overcoming action costs) (H. E. den Ouden et al., 2015; Meyniel et al., 2016) neuromodulatory systems, moreover, encode partly dissociable aspects of goal-directed actions that involve learning about costs and benefits, which together support optimal decision-making. These observations are noteworthy because the initial fight-or-flight response triggers a large-scale reorganization of brain networks that is driven by alterations in the firing mode of midbrain dopaminergic ventral tegmental area and noradrenergic *locus coeruleus* neurons (Arnsten, 2015; Hermans et al., 2014); neurons that signal prediction errors (Steinberg et al., 2013) and that are also responsive to reward value, action cost and energy expenditure (Del Arco, Park, & Moghaddam, 2020; Varazzani et al., 2015). Thus, catecholaminergic mechanisms that are recruited by the fight-or-flight response may differentially impact cost and benefit reinforcement learning, resulting in a potential scenario in which costs and benefits exert asymmetric influences on decisions.

As mentioned above, the central (i.e., neural) effects of acute stress trigger a shift in cognitive strategies, including reinforcement learning. The peripheral counterpart of the acute stress response, however, mobilizes the energy (i.e., adrenaline-mediated glucose release (de Kloet et al., 2005; Russell & Lightman, 2019)) that is required to exert effortful actions aimed at preserving homeostasis (Cannon, 1915). Therefore, decision-making and learning policies regarding physical costs may be *especially* susceptible to stress: both via computational (neural) mechanisms that support learning about and representation of action cost, as well as peripheral mechanisms that co-determine the amount of available energy that can be directed towards effortful actions. Indeed, preliminary evidence suggests that acute stress alters the willingness to exert physical effort for rewards (Bryce & Floresco, 2016) and reward-associated cues in a Pavlovian-instrumental transfer context (Pool, Brosch, Delplanque, & Sander, 2015).

How acute stress impacts reinforcement learning involving costs *and* benefits has not been investigated to date in humans. Based on the above considerations, we expect that computationally frugal learning strategies, in concert with increased energy availability, during acute stress should asymmetrically impact cost versus benefit learning. Using an acute stress-induction paradigm, a cost-benefit learning paradigm and computational model of cost-benefit reinforcement learning (Skvortsova et al., 2017; Skvortsova et al., 2014), we demonstrate that acute stress asymmetrically prioritizes reward (maximization) learning over physical effort (minimization) learning. Better benefit versus cost learning results from a stress-induced change in the influence of RPEs versus EPEs on future decisions, and is associated with altered pupil encoding of RPEs, EPEs, and subjective decision value. These results reveal how neural and peripheral mechanisms that support the fight-or-flight response may facilitate a shift in reinforcement learning strategies that confers strategic benefits during acutely stressful situations (e.g., ignoring high action costs to achieve a desirable outcome), yet might also give rise to maladaptive behaviour (e.g., stress-induced relapse in substance users).

## Results

### Experiment design

Healthy human participants were randomly assigned to the acute stress (19 males/21 females; age M=23.48, SD=3.94) or no-stress control condition (18 males/22 females; age M=23.80, SD=4.23) of the Maastricht Acute Stress Task (MAST) (Smeets et al., 2012), a validated psychological and physical stress-induction paradigm (see Materials and Methods). Immediately post-MAST and within the confines of the acute stress response (Hermans et al., 2014), all participants completed a ∼40 minute probabilistic cost-benefit reinforcement learning paradigm, adapted from Skvortsova et al. (Skvortsova et al., 2017; Skvortsova et al., 2014), in which they learned to select stimuli with high reward value (20 Eurocents) and avoid stimuli with high action cost (exerting grip force above a pre-calibrated individual threshold of 50% maximum voluntary contraction for 3000ms), followed by a surprise test phase. A detailed overview of the paradigm is provided in Figure 1 and the Materials and Methods. Pupil size was continuously recorded while participants performed the task (see Materials and Methods).

**Figure 1.**
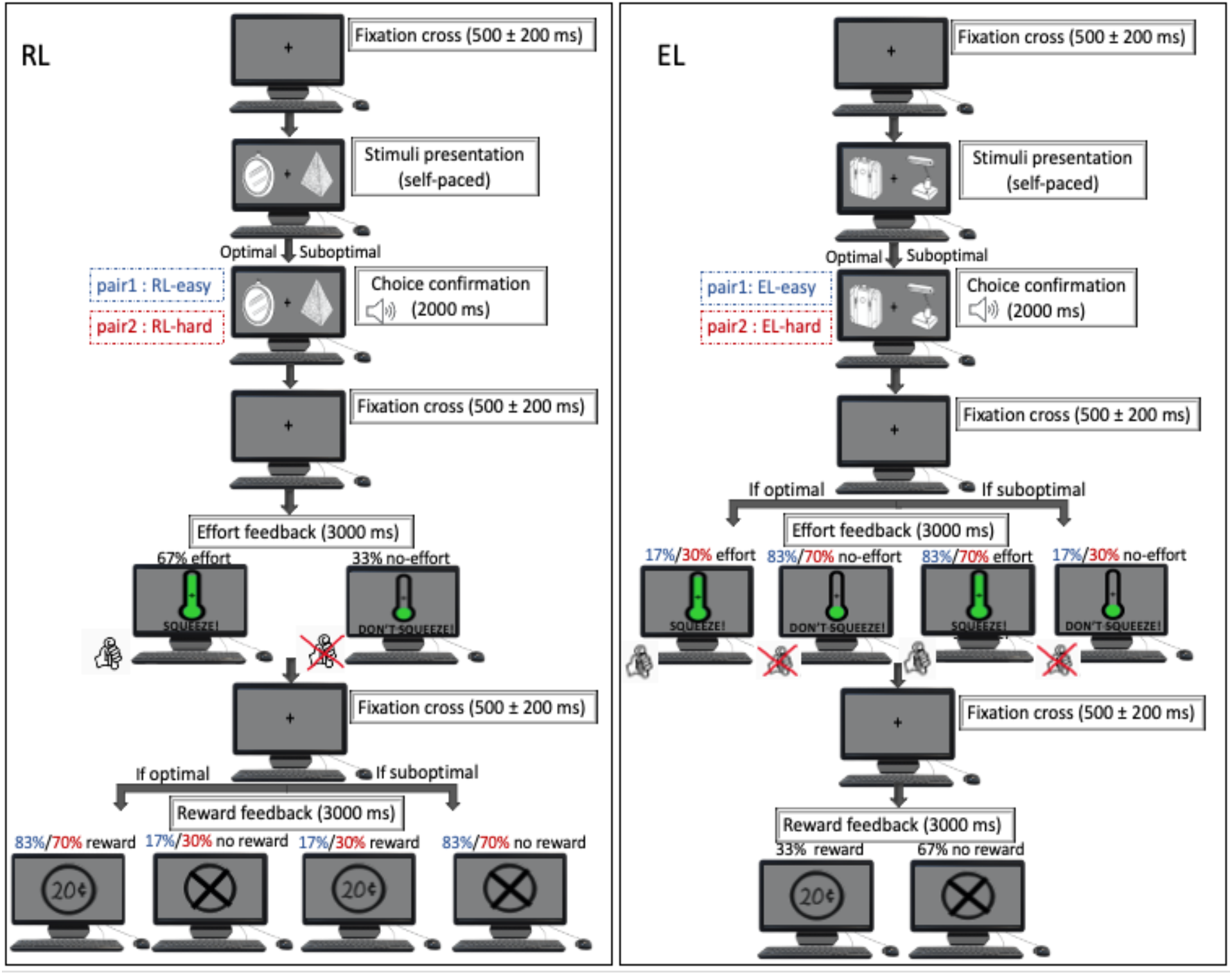
Reward maximisation/action cost minimization reinforcement learning task. Visual depiction of the learning phase. Participants were presented with four distinct stimulus pairs, and all stimuli were associated with a predetermined chance of a €0.20 monetary reward (versus no reward) and a chance of having to exert physical effort (grip force) using a dynamometer (versus no grip force required). Stimulus-outcome probabilities were yoked in such a way that, for a given pair, two stimuli *only* differed in the probability of earning a reward (“reward learning”, RL, left) or the probability of having to exert effort (“effort learning”, EL, right). That is, for RL (left)/EL (right) pairs, reward/effort outcomes were choice-dependent, respectively (see “Reward feedback” for RL and “Effort feedback” for EL for outcome contingencies). For RL pairs, effort outcomes were independent of choice and fixed (see “Effort feedback” for RL), while for EL pairs, reward outcomes were independent of choice and fixed (see “Reward feedback” for EL). Percentages in blue and red refer to outcomes for the Easy RL/EL and Hard RL/EL pair, respectively.

### Acute stress manipulation

We first ascertained whether the acute stress manipulation was successful. Subjective stress, physiological and neuroendocrine measurements are displayed in Figure 2. Acute stress and no-stress control groups did *not* differ on physiological, subjective stress, or neuroendocrine measurements pre-MAST (all *p-values*>0.05). We observed significant Condition-by-Time interactions for subjective stress ratings [PANAS negative: *F*(1,78)=52.66, *p*<0.001, *n^2^_G_*=0.10; PANAS positive: *F*(1,78)=9.82 *p*=0.002, *n^2^_G_*=0.02] and physiological measures [systolic blood pressure (SBP): *F*(1,78)=15.50, *p*<0.001, *n^2^_G_*= 0.04; heart rate: *F*(1,78)=6.83, *p*=0.011, *n^2^_G_*= 0.02]. Simple main effect analyses revealed that only the acute stress group exhibited pre-to-post *increases* in negative affect [control pre-post: *t*(39)=4.21, *p*<0.001; stress pre-post: *t*(39)=-6.17, *p*<0.001; control-stress post-MAST: *t*(55.1)=-5.78, *p*<0.001], and greater pre-to-post *decreases* in positive affect [control pre-post: *t*(39)=4.09, *p*<0.001; stress pre-post: *t*(39)=6.45, *p*<0.001; control-stress post-MAST: *t*(72.8)=2.53, *p*=0.014] (Figure 2). Similarly, only the acute stress group exhibited stress-induced *increases* in SBP [control pre-post: *t*(39)=1.60, *p*<0.117; stress pre-post: *t*(39)=-3.66, *p*<0.001; control-stress post-MAST: *t*(69.1)=-3.27, *p*=0.002] and heart rate [control pre-post: *t*(39)=1.21, *p*=0.234; stress pre-post: *t*(39)=-2.78, *p*=0.008; control-stress post-MAST: *t*(76.9)=-3.14, *p*=0.002] (Figure 2a-d).

**Figure 2.**
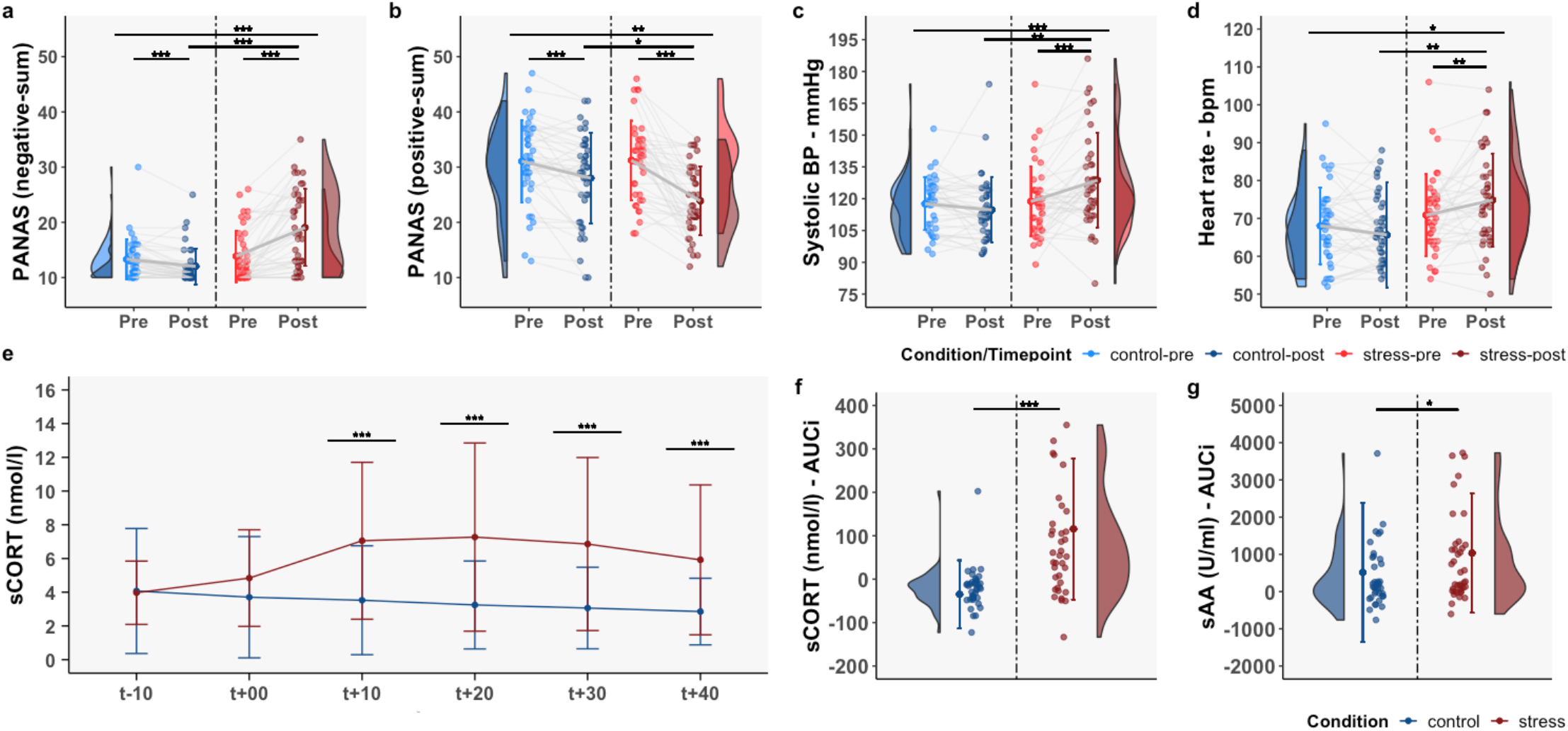
Neuroendocrine, physiological and subjective stress ratings. PANAS negative (**a**) and positive (**b**) subscale sum scores, systolic blood pressure (mmHg: millimetres of mercury; **c**) and heart rate (bpm: beats per minute; **d**) are displayed for no-stress control (blue) and acute stress (red) groups separately for pre (light blue/red) and post (dark blue/red) MAST time points. SCORT responses for both conditions across the 6 timepoints are displayed in panel **e** (“t_+00_” represents the first post-MAST measurement, and the start of the reward maximization/action cost reinforcement learning paradigm; “t_-10_” represent a baseline sample). Panel **f** and **g** show AUCi for sCORT (nmol/l: nanomoles per litre) and sAA (U/mL: Units per millilitre) responses for both MAST conditions. Significant differences are denoted by asterisks (*: *p* < 0.05, **: *p* < 0.01, ***: *p* < 0.001). In the upper panel, the top line denotes a significant Condition-by-Time interaction; lower lines represent simple main effects of Condition or Time.

An expected Condition-by-Time interaction was found for salivary cortisol (sCORT) responses [*F*(5,390)=18.05, *p*<0.001, *n^2^_G_*= 0.04], with the acute stress group displaying greater sCORT levels 10 min post-MAST and onwards (all *p-values*<0.01). We additionally observed a main effect of Condition on sCORT area-under-the-curve with respect to increase: (AUCi) (Pruessner, Kirschbaum, Meinlschmid, & Hellhammer, 2003) (*t*(56.32)=-5.28, *p*<0.001) and salivary alpha-amylase (sAA) AUCi (*t*(67.45)=-2.50, *p*=0.015; after excluding one extreme outlier from the control group), suggesting greater sCORT and sAA levels in response to acute stress (Figure 2e-g). These results confirm that the MAST robustly induced stress on all levels of inquiry.

### Participants use reinforcement learning to optimize decisions

Next, we investigated whether participants in both conditions exhibited evidence of reinforcement learning to optimize actions, which in this paradigm should be reflected by an increased tendency to select stimuli with high reward value and avoid stimuli with high action cost as a function of increasing number of stimulus pair presentations (i.e., “time”). This intuition was confirmed by a main effect of Time on two distinct trial types: trials on which participants could learn to accumulate frequent rewards while the probability of effort (action cost) was kept constant (RL; selecting the stimulus more frequently associated with €0.20) [control: *F*(2,78)=10.16, *p*<0.001, *n^2^_G_*= 0.06; stress: *F*(2,78)=20.44, *p*<0.001, *n^2^_G_*= 0.17] and trials on which participants could learn to frequently avoid effort while the probability of reward was kept constant (EL; selecting the stimulus more frequently associated with avoidance of physical energy expenditure) [control: *F*(2,78)=12.35, *p*<0.001, *n^2^_G_*= 0.07; stress: *F*(2,78)=9.76, *p*<0.001, *n^2^_G_*= 0.05]. We additionally observed greater than chance-level performance (≥0.5) on both trial types during the final part of the task (presentation 21-30; all *p*-values<0.001 Figure Supplement 1).

### Asymmetric cost-benefit reinforcement learning during acute stress

After having observed evidence for reward (maximization) learning and action cost (minimization) learning, we tested our key assumption; that acute stress would induce a reprioritization in learning to maximize reward value versus learning to minimize action cost. Crucially, we observed a significant Condition-by-Trial Type interaction [*F*(1,78)=6.53, *p*=0.013, *n^2^_G_*= 0.039] (Figure 3a) with pairwise comparisons indicating that the acute stress group performed significantly better on RL than EL trials [*t*(39)=5.40, *p*<0.001], while the no-stress control group performed similarly on both trial types [*t*(39)=1.01, *p*=0.320]. A main effect of Condition on RL-EL accuracy difference scores [*t*(74.02)=-2.55, *p*=0.013] (Figure 3b) and one-sample t-tests revealed that RL-EL accuracy difference scores were significantly greater than zero in the acute stress group but not in the no-stress controls. Simple main effects of Condition on RL [*t*(65.9)=-1.75, *p*=0.085] and EL [*t*(77.5)=1.80, *p*=0.076] performance showed numerical trends for group differences that failed to reach significance.

**Figure 3.**
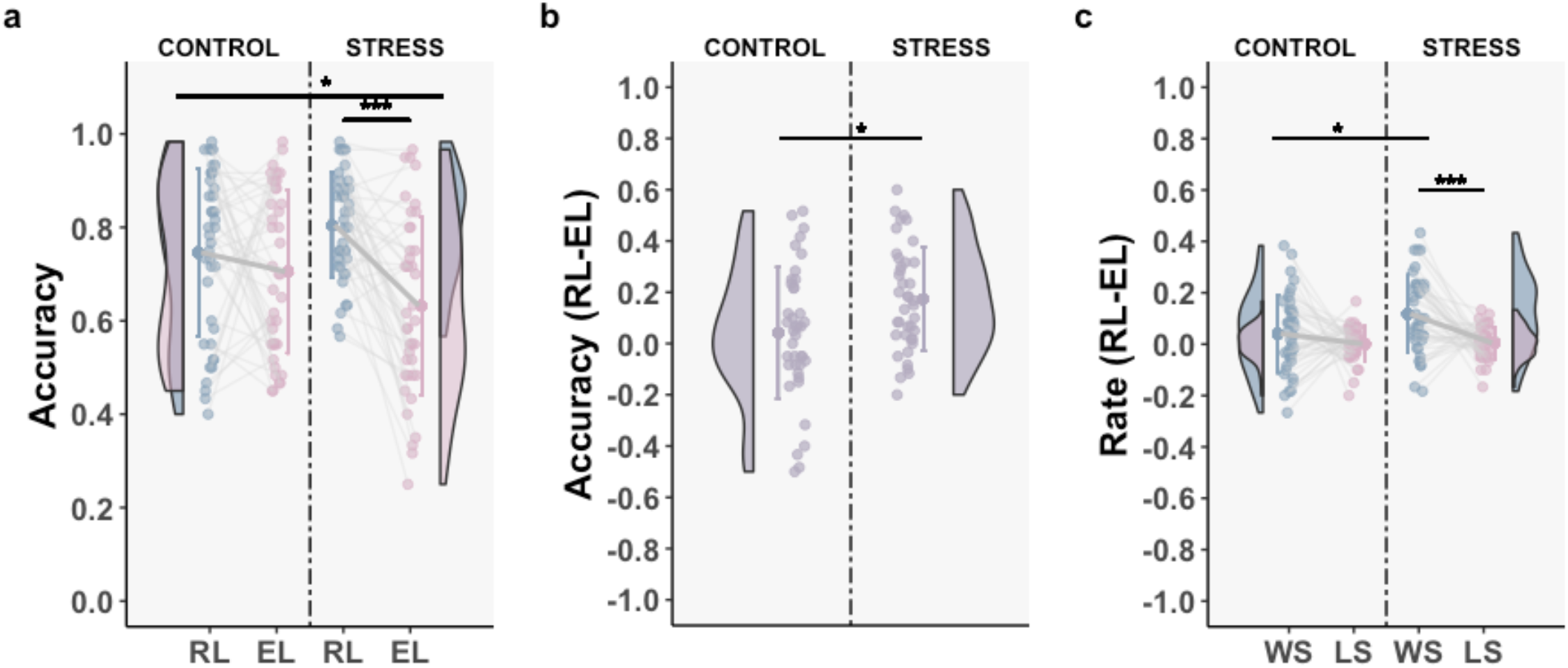
Acute stress leads to improved benefit versus cost learning. Panel **a**: Average accuracy (choices of the optimal stimulus) for RL and EL trials, for each condition separately. Panel **b**: RL-EL average accuracy difference scores. Panel **c:** Win-stay (WS_RL_-WS_EL_) and lose-shift (LS_RL_-LS_EL_) difference scores for each condition separately. Means ± SD, individual data points, distribution and frequency of the data are displayed. In panel **a**, the top line indicates a significant Condition-by-Trial type interaction. Significant differences are denoted by asterisks (*: *p* < 0.05, **: *p* < 0.01, ***: *p* < 0.001). Source files of task performance data used for the analyses are available in the Figure 3 – Source Data 1.

When we included participants that still performed at chance level at the end of the learning phase (see Participants) in the Condition-by-Trial Type interaction analysis, the interaction remained significant [*F*(1,91)=7.30, *p*=0.035, *n^2^_G_*= 0.04], with the acute stress group displaying better RL vs. EL performance (*t*(46)=5.83, *p*<0.001), while no-stress controls performed similarly on both trial types (*t*(45)=1.24, *p*=0.22). Participants in the acute stress group outperformed participants in the no-stress control group on RL (*t*(91)=2.04, *p*=0.04), but no simple main effect of Condition was observed for EL (*t*(91)=-1.67, *p*=0.1). These participants were excluded for all other analyses reported below.

The use of different reinforcement probabilities for each RL and EL pair (Figure 1) allowed us to discern whether the observed pattern of results reflected a *specific* change in cost (EL) versus benefit (RL) reinforcement learning, or a more general impairment in reinforcement learning for more difficult stimulus-response associations. We found no evidence for the latter scenario in Condition-by-Trial Type-by-Difficulty [*F*(1,78)=1.05, *p*=0.310] or Condition-by-Difficulty [*F*(1,78)=0.36, *p*=0.549; Figure Supplement 2] interaction analyses.

When we investigated the use of win-stay (resampling a stimulus following a positive outcome) and lose-shift (switching to the other stimulus following a negative outcome) strategies (Hanneke E. M. den Ouden et al., 2013), we observed no significant Condition-by-Strategy interaction [*F*(1,78)=2.99, *p*=0.087, *n^2^_G_*= 0.03]. However, *post hoc* comparisons revealed that participants in the acute stress condition [*t*(39)=-3.73, *p*<0.001] but not those in the no-stress control condition [*t*(39)=-1.30, *p*=0.200], exhibited different win-stay compared to lose-shift (difference) rates. Separate Condition (main effect) analyses indicated that acute stress participants compared to no-stress controls were more likely to win-stay for rewards (RL trials) than for avoidance of action cost (EL trials) [*t*(78)=-2.28, *p*=0.025], but not for lose-shifting for reward omissions (RL trials) compared to exerting effort (EL trials) [*t*(77.7)=-0.23, *p*=0.820]. Differences in win-stay rates for RL compared to EL trials (one-sample t-test) were greater than zero for the acute stress [*t*(39)=4.91, *p*<0.001] but not the no-stress control group [*t*(39)=1.68, *p*=0.101] (Figure 3c).

Taken together, our model-free results indicate that acute stress leads to a reinforcement learning strategy that favours learning to maximise reward value over minimisation of action cost, which based on analyses of win-stay/lose-shift rates, could be attributed to increased sensitivity to positive reinforcement (i.e., reward delivery) compared to negative reinforcement (i.e., avoidance of physical effort).

### Asymmetric cost-benefit reinforcement learning biases actions in acute stress subjects

During a surprise 64-trial test phase (D. Hernaus, Gold, Waltz, & Frank, 2018), we asked participants to discriminate original and novel combinations of stimuli on the basis of reward value or action cost without receiving feedback (n=16 trials for original combinations; n=48 for novel combinations; see Materials and Methods). The surprise test phase allowed us to assess learned choice tendencies without having to arbitrarily choose a given number of final learning phase trials, during which participants may still learn. This approach also allowed us to assess the degree to which learned tendencies would carry over to novel contexts.

First, both groups chose the optimal (most rewarding/effort avoiding) stimulus on surprise test phase trials involving the original four pairs [one-sample t-test against chance; control_RL_: *t*(39)=8.73, *p*<0.001; control_EL_: *t*(39)=3.72, *p*=0.002; stress_RL_: *t*(39)=13.54, *p*<0.001; stress_EL_: *t*(39)=4.47, *p*<0.001], confirming that both groups had developed a preference for the optimal stimulus.

Although we observed no Condition-by-Trial type (reward value, action cost discrimination) interaction or main effects of Condition for novel stimulus combinations [*F*(1,78)=1.10, *p*=0.298, *n^2^_G_*= 0.01; stress vs. controls reward value discrimination: *t*(75.9)= 0.15, *p*=0.878; stress vs. controls action cost discrimination: *t*(78)= 1.77, *p*=0.080], pairwise comparisons revealed that the acute stress group performed better on reward discrimination compared to action cost discrimination trials [*t*(39)=-2.23, *p*=0.032], while no-stress controls performed similarly on both trial types [*t*(39)=-0.87, *p*=0.387]. These results provide some evidence that a reward maximisation-over-action cost minimisation reinforcement learning policy might bias future actions in novel contexts (Figure Supplement 3).

### Computational cost-benefit reinforcement learning model: Model Fitting, Selection, Demonstrations, and Simulations

To uncover latent mechanisms by which acute stress affects cost-benefit reinforcement learning, we turned to computational cognitive modelling. Trial-by-trial choices of participants were fit to all candidate models described in the Materials and Methods (see Model Space). To calculate model fit, log likelihood was updated trial-wise by the log of the probability of the observed choice, calculated via a softmax rule (see Materials and Methods, equation 6), and best-fitting parameters were identified using fmincon in MATLAB v.2019B (Mathworks, Natick, MA, USA).

Bayesian Model Selection (BMS; spm_BMS function in SPM12, http://www.fil.ion.ucl.ac.uk/spm/software/spm12/) using the Akaike Information Criterion (AIC) as a fit statistic that penalizes for the number of model parameters (Myung, Tang, & Pitt, 2009), suggested that the 2LR_γ model was the most likely model, as indicated by the protected exceedance probability (pxp, φ = 0.99) (Rigoux, Stephan, Friston, & Daunizeau, 2014) and expectation of the posterior [p(r|y), 0.70] (Figure 4a for p(r|y) of all candidate models). We note that 2LR_γ remained the most likely model when we considered additional models with greater redundancy and/or lesser biological plausibility (e.g., models with all combinations of reward value/action cost discounting *and* weight parameters).

**Figure 4.**
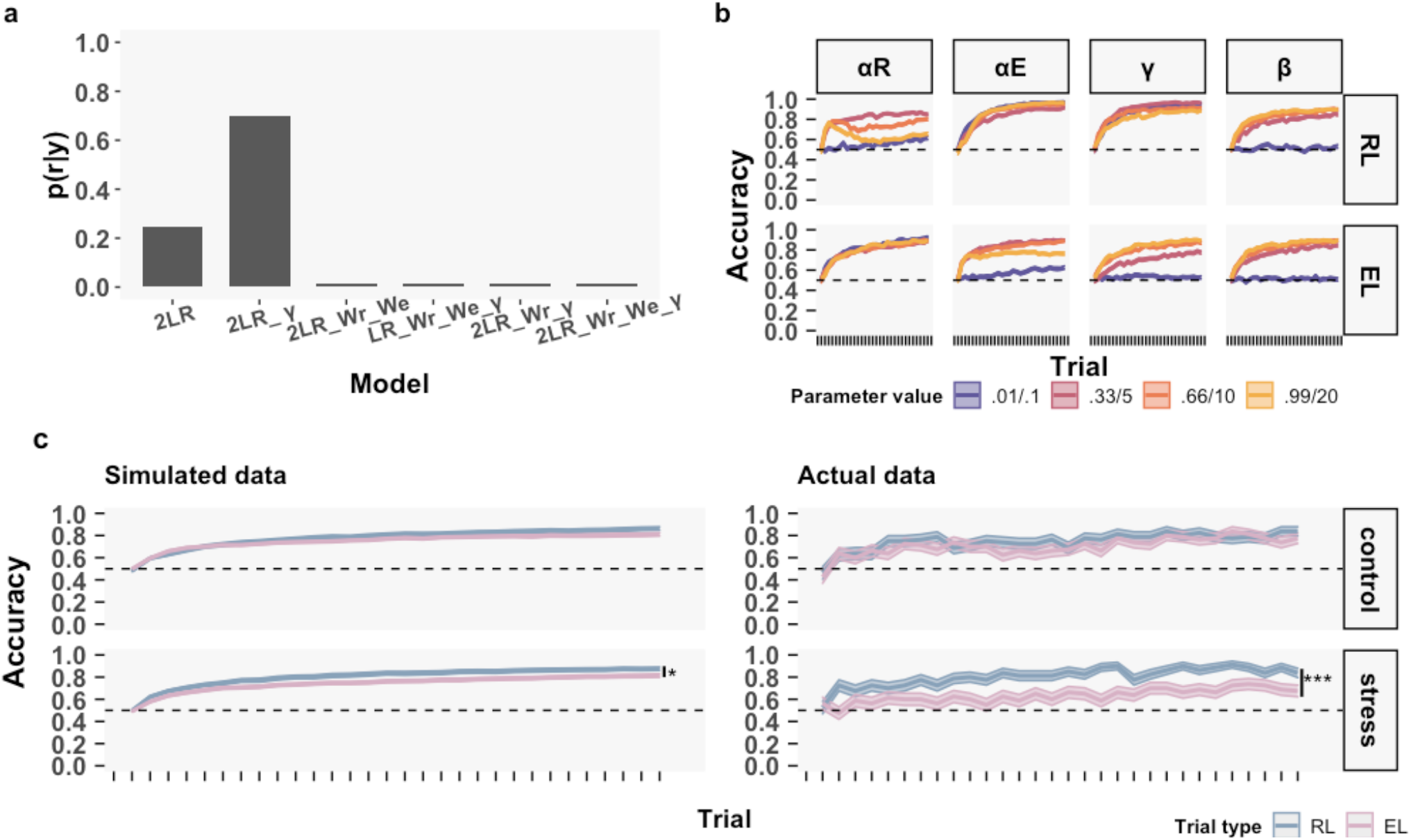
Model selection, demonstrations, and *post hoc* simulations of the winning model. Panel **a**: Expectation of the posterior for all candidate models. Panel **b**: Model demonstrations. To demonstrate how different parameter values within the 2LR_γ architecture impact choice preferences for the optimal stimulus (“accuracy”), α_R_, α_E_, and γ were set to 0.01/0.33/0.66/0.99, while β, a non-linear parameter, was set to 0.1/5/10/20. Parameter effects were always demonstrated for a single parameter (columns), while all other parameter values were kept constant (α_R_ and α_E_=0.25, γ=1, β=25). Greater values of α_R_ and α_Ε_ selectively increase the speed with which the agent develops a preference for the optimal RL and EL stimulus, respectively. Lower values of γ produce an asymmetric decision-making policy that emphasises reward value over action cost, leading to better performance on RL versus EL trials, while greater values of γ correct this asymmetric choice bias. Finally, greater β values lead to more deterministic sampling of optimal stimuli. Panel **c**: Post-hoc simulations after fixing β and γ to group-level averages. Coloured lines represent mean ± SD. Dashed lines denote chance level (0.5). *: *p* < 0.05, **:*p* < 0.01, ***: *p* < 0.001.

The 2LR_γ model contains separate learning rates that weight the importance of RPEs and EPEs (α_R_, α_E_), an action cost discounting parameter (γ), and an inverse temperature parameter (β), which in previous work could account for performance on a conceptually similar cost-benefit learning task (Skvortsova et al., 2014). To demonstrate the effect of changes in parameters values on choice preferences within the 2LR_γ architecture, we first simulated choices from 50 artificial agents (averaged across 10 repetitions) performing the reward maximization/action cost minimization reinforcement learning task using a range of parameter values. As expected, greater values of α_R_ and α_E_ primarily impacted the speed of RL and EL choice preferences, while low values of γ lead to asymmetric choice preferences through discounting of action cost, and lower values of β lead to non-selective increases in random sampling (Figure 4b).

In *post hoc* simulations, i.e., generating participant choices using the obtained parameters, we additionally observed moderate-to-high correlations between simulated and empirical RL/EL for the acute stress and no-stress control group [*ρ*_RL_control_ = 0.55, *p* < 0.01; *ρ*_RL_stress_ = 0.84, *p* < 0.01; *ρ*_EL_control_ = 0.56, *p* < 0.01; *ρ*_EL_stress_ = 0.77, *p* < 0.01; see Figure Supplement 4], although the canonical performance difference in RL versus EL accuracy was not selective to the acute stress group [*t_control_*(39)=-6.72, *p*<0.001; *t_stress_*(39)=-6.01, *p*<0.001]. However, after we fixed β and γ to group-level averages, to better demonstrate the effect of group differences in the learning rate parameters, we recovered a small but significant simulated difference in RL versus EL performance for the acute stress group [*t*(39)=2.27, *p*=0.029], which was not predicted in the no-stress control group [*t*(39)=0.91, *p*=0.367] (Condition-by-Trial Type interaction: [*F*(1,78)= 0.77, *p*=0.38, *n^2^_G_*= 0.006]) (Figure 4c for empirical versus simulated data, averaged across 100 repetitions per subject).

Importantly, even if a given model is the most likely one based on model fitting and *post hoc* simulation results from the entire sample, there is still the possibility that different models can better explain task performance in the no-stress control and acute stress condition. When repeating BMS for each condition separately, 2LR_γ was the most likely model in the no-stress control group [φ=0.99, p(r|y)=0.83], while for acute stress subjects 2LR_γ was not convincingly the most likely model [φ=0.47 p(r|y)=0.46]. Here, the 2LR model (containing α_R_, α_E_, and β parameters) was equally likely to be the optimal model [φ=0.53 p(r|y)=0.47]. Post-hoc simulations from the 2LR model also correlated with actual data, both for no-stress control [*ρ*_RL_ = 0.65, *p* < 0.001; *ρ*_EL_ = 0.60, *p* < 0.001] and acute stress participants [*ρ*_RL_ = 0.81, *p* < 0.001; *ρ*_EL_ = 0.75, *p* < 0.001].

Similar to the 2LR_ γ model (results discussed in next section), the 2LR model seemingly also explained stress-induced changes in cost-benefit reinforcement learning via changes in learning rates; in the 2LR model, the acute stress group exhibited greater values of α_R_ versus α_E_ (t(39)=2.65, *p*=0.01), while no-stress control subjects did not (*t(*39_)_=0.69, *p*=0.50) (Condition-by-Learning Rate interaction: [*F*(1,78)=2.88, *p*=0.094, *n^2^_G_*= 0.01]. The difference in learning rates between 2LR_γ (where α_R_ and α_E_ are similar for the acute stress group, see next section) and 2LR (where α_R_ > α_E_ for the acute stress group) can be explained by the absence of discounting parameter γ: 2LR is a special case of 2LR_γ, where γ=1, and thus asymmetric effects of acute stress on reward value maximization and action cost minimization can only be explained by *dissimilarity* in learning rates.

Although the effects of acute stress on reward value and action cost learning rates are opposite in 2LR_ γ versus 2LR architectures, these results bolster our confidence in the overall model space, as well as the interpretation that acute stress primarily impacts reward value and action cost learning rates, and *not* discounting. The observations that I) 2LR_ γ fit better in the entire group of participants, II) 2LR is fully contained within the 2LR_ γ model, and III) 2LR_ γ displayed good recoverability (see *below)* motivated our choice to focus on the 2LR_γ model.

In model recoverability analyses i.e., re-fitting the simulated data from the model to all candidate models (Wilson & Collins, 2019), BMS confirmed that the simulated 2LR_γ data (that is, simulations *without* fixed parameters) were most likely to be generated from 2LR_γ [φ=0.99, p(r|y)=0.71].

To assess the stability of 2LR_γ parameters, we repeated model fitting using a Bayesian hierarchical model fitting approach consisting of two steps, as described previously (Daw, 2011; Frey, Frank, & McCabe, 2019). In the first step we fit the 2LR_γ model to trial-wise choices to obtain subject-specific parameters; in a second step we again fit the model to trial-wise choices, but this time we used the group-level average and covariance matrix of every parameter as priors, thereby shrinking the parameter search space. Motivated by recent work showing that group-specific priors, compared to a single prior for the entire sample, can better account for between-group differences in task performance, as well as improve parameter robustness and recoverability (Valton, Wise, & Robinson, 2020), we used separate mean and covariance matrices for the acute stress and no-stress control groups.

Highly similar parameter estimates were obtained after hierarchical fitting (for parameter estimates after Bayesian hierarchical model fitting see Figure Supplement 5). Similar to *post hoc* simulations using parameters from the non-hierarchically fit 2LR_γ model, we observed moderate-to-high correlations between empirical and simulated data using parameters obtained from the hierarchically fit model [*ρ*_RL_control_ = 0.65, *p* < 0.01; *ρ*_RL_stress_ = 0.84, *p* < 0.01; *ρ*_EL_control_ = 0.37, *p* =0.02; *ρ*_EL_stress_ = 0.78, *p* < 0.01; see Figure Supplement 6]. All in all, these results confirm parameter stability within the 2LR_γ architecture.

In light of model fitting results, post-hoc simulations, model and parameter recoverability analyses, we used parameters and trial-by-trial predictions of the non-hierarchically fit 2LR_ γ model in all analyses reported below.

### Acute stress selectively reduces the difference between reward and action cost learning rates

Comparing 2LR_γ parameters between conditions, we observed a significant Condition-by-Learning Rate (α_R_, α_E_) interaction [*F*(1,78)= 6.42, *p*=0.01, *n^2^_G_*= 0.03; 95% highest density interval (HDI) for Bayesian mixed ANOVA = −0.405 to −0.023, mean= −0.219]. with greater EPE relative to RPE learning rates in no-stress control participants [*t*(39)=-4.75, *p*<0.001], while learning rates in the acute stress group did not significantly differ [*t*(39)=-1.61, *p*=0.116]. No between-group differences in α_R_ and α_E_ or in the other parameters (γ, β) were observed (all *p-values*>0.05) (Figure 5).

**Figure 5.**
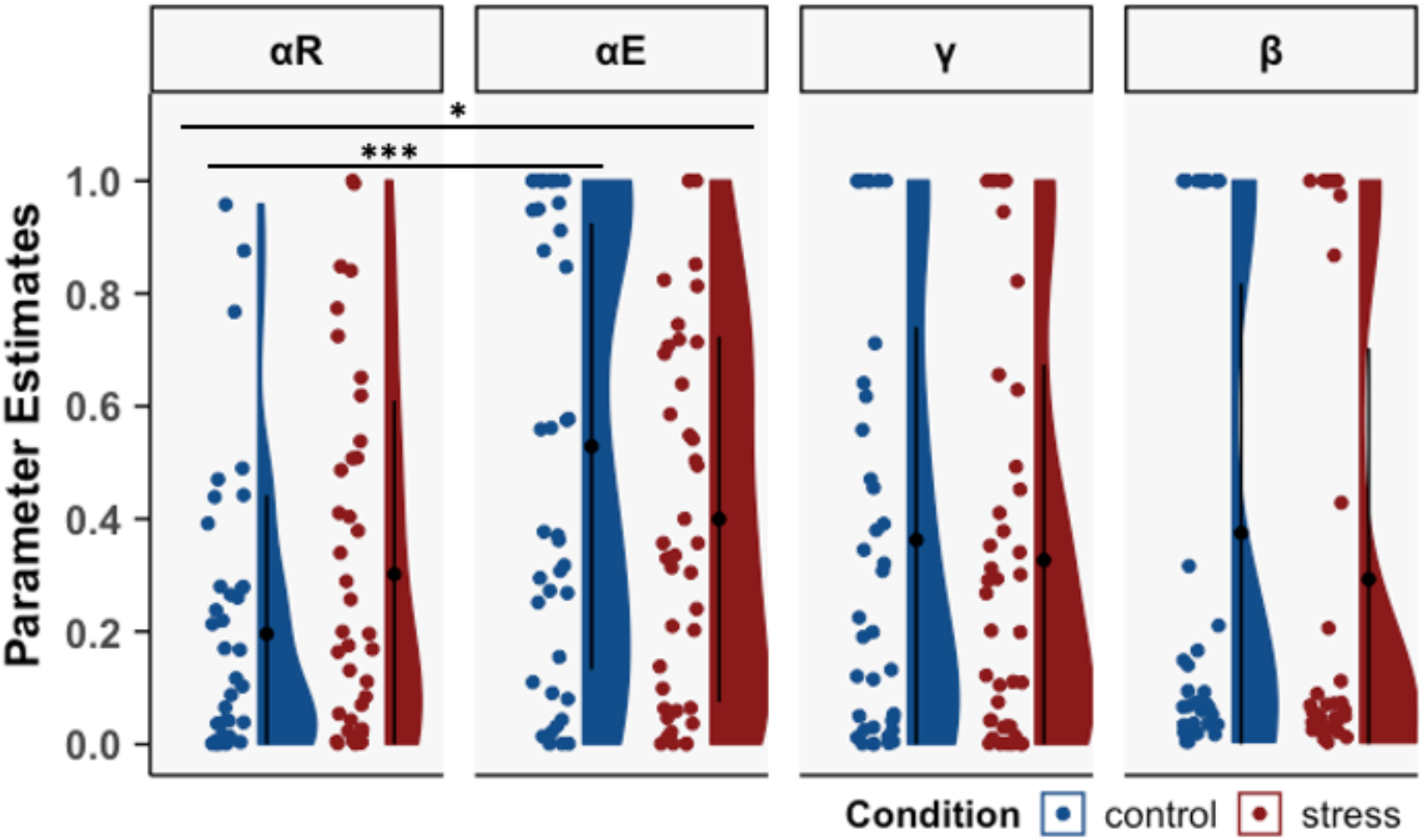
Acute stress reduces the difference between reward and effort prediction error learning rates. Free parameters (αR, αE, γ, β) of the winning 2LR_ γ model for both groups. Black lines denote means ± SD, dots represent individual data points, and the violin-like shape denotes distribution and frequency of the data. *: *p* < 0.05, **: *p* < 0.01, ***: *p* < 0.001.

Paradoxically, symmetric reward value and action cost learning rates in the presence of lower values of γ will lead to more efficient RL compared to EL. This is because lower values of γ bias decisions towards reward value (via *greater* discounting of action cost) and similar absolute values of α_R_/α_E_ will not counteract this bias. *Asymmetric* learning rates (α_E_>α_R_) in combination with lower values of γ, however, will lead to more symmetric performance on RL and EL trials via more efficient updating of action cost versus reward expectations. This interpretation is supported by our demonstration of model parameters (Figure 4b) and *post hoc* simulations (Figure 4c), as well as the observation that lower values of γ (i.e., greater action cost discounting) were associated with greater learning rate asymmetry (α_E_>α_R;_ more efficient EL) in no-stress controls (ρ=-0.40, *p*=0.040), who displayed similar RL and EL performance. These results demonstrate that, in a context where all decisions involve a potential cost and benefit, acute stress selectively reduces the difference between EPE and RPE learning rates, while leaving action cost discounting and choice stochasticity unaffected. The direction of the change in learning rates (i.e., greater similarity) implies a stress-induced failure to modulate learning rates in the service of overcoming an asymmetric choice bias that emphasises reward value.

In analyses using posterior parameters obtained from the hierarchically fit model, we recovered the key Condition-by-Learning Rate interaction (95% HDI for Bayesian mixed ANOVA = −0.406 to −0.128, mean=-0.269) [and acute stress and no-stress control subjects differed from each other on α_E_ (95% HDI = 0.0841 to 0.281, mean=0.183) but not α_R_ (95% HDI = −0.186 to 0.0102, mean=-0.0872)] (Figure Supplement 7). Similar to the non-hierarchically fit parameters, acute stress and control subjects did not differ on posterior estimates of γ (95% HDI = −0.0914 to 0.139, mean=0.0182) and β (95% HDI = −0.0331 to 0.213, mean=0.0895).

### Pupil size fluctuations track asymmetric cost-benefit reinforcement learning during acute stress

We employed pupillometry to better understand whether task-relevant computational processes may be encoded by fluctuations in pupil dilation, which are thought to be controlled by ascending midbrain modulatory systems that play a role in value-based decision-making and the acute stress response (Arnsten, 2015; Hermans et al., 2011; Joshi, Li, Kalwani, & Gold, 2016).

In model-free analyses – that is, comparing bins of pupillometry data between conditions - we observed no main effect of Condition on pupil size fluctuations during choice, effort outcome, and reward outcome epochs, suggesting that acute stress was not associated with more general changes in pupil size (all bin-level *p*>0.05; Figure 6, a. Model-free; Figure Supplement 8 for all effort trials).

**Figure 6.**
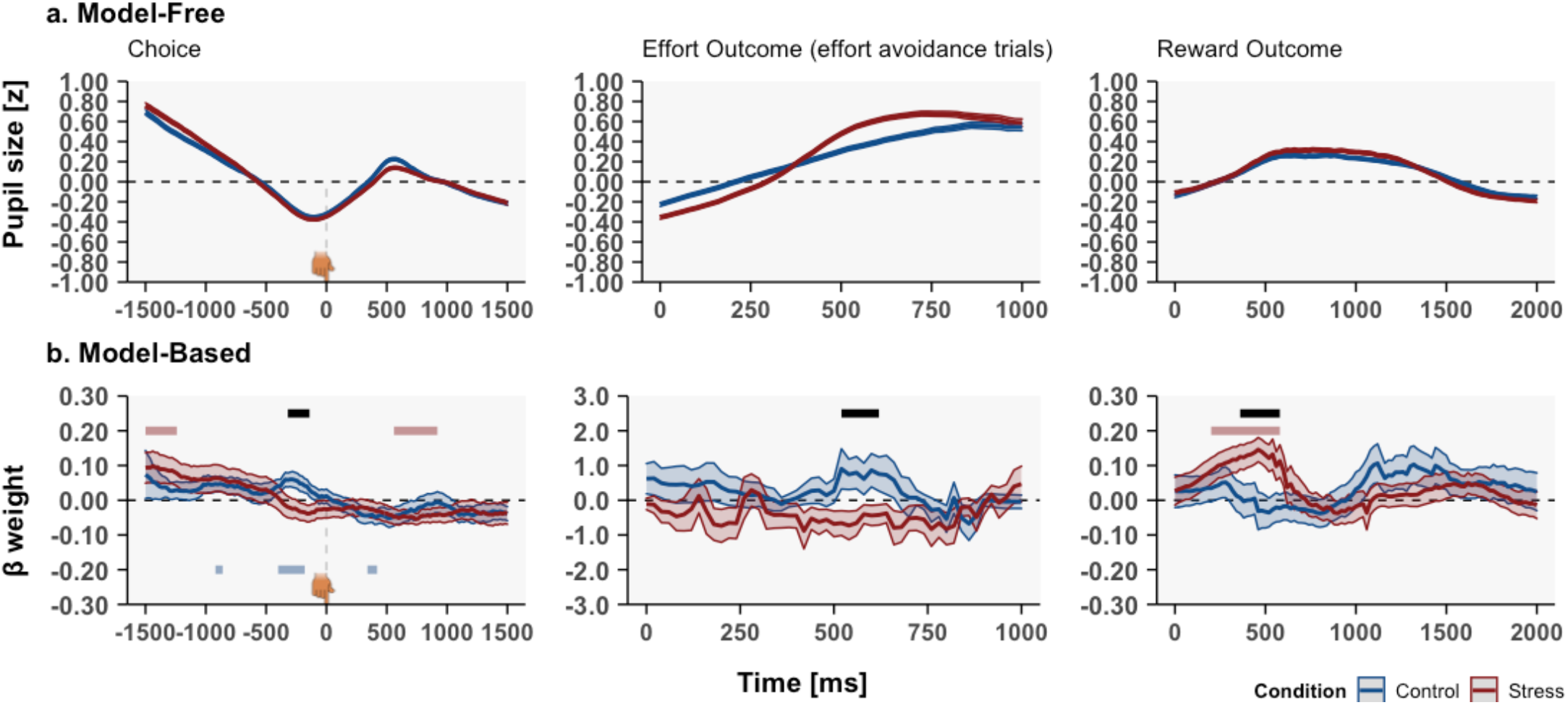
Model-based analysis reveals altered pupil encoding of prediction errors and decision value during acute stress. **a**. Model-free analyses of pupil size during choice, effort outcome, and reward outcome phase revealed no main effect of Condition (no-stress control, acute stress). **b.** Model-based analyses revealed a stress-induced shift in pupil encoding of subjective decision value (left), action cost prediction errors (middle) and reward prediction errors (right). Black line indicates significant main effect of Condition; blue and red line indicate significance against zero for no-stress control and acute stress groups, respectively (cluster and bin level *α*_permute_<0.05, 2000 permutations). Group differences in pupil encoding of action cost and reward prediction errors were observed at similar times (note the x-axis differences for effort outcome and reward outcome epochs). Source files of pupillometry data used for the analyses are available in Figure 6 – Source Data 2.

Next, we conducted model-based pupillometry analyses (Lawson, Bisby, Nord, Burgess, & Rees, 2020) to understand how trial-wise estimates of computational processes of interest were encoded by fluctuations in pupil size. These analyses revealed effects of Condition on pupil encoding of subjective decision value, EPEs and RPEs (Figure 6, b. Model-based) in a manner commensurate with task performance results. First, immediately prior to the stimulus choice, acute stress reduced pupil encoding of subjective decision value, as evidenced by the absence of an association between pupil size and subjective decision value (control>stress; stress n.s., control>0). Second, briefly after the presentation of effort avoidance outcomes, both groups exhibited different pupil size-EPE associations, with no-stress controls showing a non-significant numerically positive association between pupil size and action cost prediction errors (control>stress, both groups n.s. different from 0). In model-based analyses using all effort outcome trials, we were not able to uncover group differences in pupil encoding of EPEs, which were likely eclipsed by prominent grip force-related effects on pupil size (Figure Supplement 8). Third, during the reward outcome phase, acute stress participants exhibited greater positive associations between pupil size and RPEs compared to no-stress controls (stress>control, stress>0, control n.s.). The average pupil size-RPE slope for bins in which no-stress control and acute stress participants differed (Figure 6b) correlated significantly with stress-induced changes in SBP [*ρ_stress_*(38)= −0.41, *p(permutation)*=0.019] and PANAS negative affect changes [*ρ_s_*_tress_= −0.46, *p(permutation)*=0.005] in the stress group.

Crucially, group differences in pupil encoding of subjective decision value, EPEs, and RPEs imply that the ascending neuromodulatory systems may have facilitated a stress-induced shift in asymmetric cost versus benefit learning.

## Discussion

Stress-induced alterations in adaptive decision-making are commonly studied using paradigms that isolate positive and negative reinforcement, such as the receipt of a reward or avoidance of a loss. However, it remains poorly understood how acute stress affects the complex process that entails learning about costs *and* benefits, a critical and pervasive feature of everyday decisions. Participants completed a paradigm in which all actions (stimulus choices) contained a potential cost (exerting physical effort) and a financial benefit (€0.20). Crucially, acute stress induced a shift in reinforcement learning strategies that improved maximization of monetary rewards relative to minimisation of energy expenditure. When presented with novel stimulus arrangements and in the absence of feedback, individuals in the acute stress condition, moreover, exhibited better discrimination of stimulus reward value compared to action cost.

Relative improvements in reward versus action cost learning align well with previous reports of enhanced reward learning during acute stress (Byrne, Cornwall, & Worthy, 2019; Lighthall et al., 2013; Petzold et al., 2010), although such effects may depend on stressor timing (Joëls, Pu, Wiegert, Oitzl, & Krugers, 2006), stressor type (Carvalheiro et al., 2020), and/or sample characteristics (Evans & Hampson, 2015; Morris & Rottenberg, 2015). While reports on action cost learning during acute stress are scarce, acute exposure to stress in rodents impairs cost-benefit decisions via a selective change in sensitivity to physical effort, a process mediated by corticotropin-releasing factor and dopamine (Bryce & Floresco, 2016). Our analyses of win-stay/lose-shift rates indicate that asymmetric cost-benefit learning can be driven by a relative increase in the sensitivity to monetary gains compared to the avoidance of costly deterrents.

How might maximization of reward value take precedence over minimisation of action cost? Acute stress leads to a redistribution of finite cognitive resources (Hermans et al., 2014): this process limits the availability of computationally intensive strategies, including working memory (Otto, Raio, Chiang, Phelps, & Daw, 2013; Qin, Hermans, van Marle, Luo, & Fernández, 2009) and goal-directed instrumental actions (Lars Schwabe & Wolf, 2011). Assuming that acute stress does not merely increase random responding - which we verified via the choice stochasticity model parameter - a computationally cheap heuristic in our task should present itself as better learning for one modality over the other. Increased energy availability (Hermans et al., 2014), insensitivity to aversive stimuli (Timmers et al., 2018), and impaired aversive value updating (Raio et al., 2017) under stress may have reduced the ability – or urgency – to dedicate cognitive resources to strategies that minimize action cost. Importantly, effort expenditure increases the perceived value of rewards (Hernandez Lallement et al., 2014; Inzlicht, Shenhav, & Olivola, 2018). Thus, frequent expenditure of physical effort, due to suboptimal action cost learning, may increase the perceived value of rewards, and thus tilt learning towards the maximization of reward value.

Using a computational model of reinforcement learning (Skvortsova et al., 2017; Skvortsova et al., 2014), we confirmed that biased cost-benefit learning can arise when inappropriate (i.e. more similar) importance is afforded to teaching signals that convey information about reward value (RPEs) and action cost (EPEs). Humans display presumably instinctive biases, such as more efficient learning from better-than-expected outcomes (Lefebvre, Lebreton, Meyniel, Bourgeois-Gironde, & Palminteri, 2017) (compared to worse-than-expected outcomes) and asymmetric “Go”/approach learning (Guitart-Masip et al., 2012) (compared to “No-Go”/avoidance learning), the latter being a bias that is also modulated by acute stress (de Berker et al., 2016). From this perspective, no-stress controls, who assigned greater importance to EPEs than RPEs, may have used a computationally costly learning strategy that provides counterweight to a decision-making policy that is biased towards the reward value of actions (captured by action cost discounting parameter γ). Paradoxically, when decisions are by default tilted towards reward value, similar reward and action cost learning rates will facilitate reward learning but hamper action cost learning. Reduced learning rate asymmetry in the presence of action cost discounting may therefore represent a computational reformulation of a heuristic that is employed when cognitively demanding learning strategies are unavailable and the policy towards energy expenditure is more liberal, such as during acute stress.

Importantly, stress-induced changes in task performance may crucially depend on the release of catecholamines in neural circuits that support motivation and learning. Dopamine’s actions at D1 and D2 receptors in the basal ganglia mediate approach and avoidance learning (Frank, Seeberger, & Reilly, 2004), and acute stress can improve associative learning by augmenting reward-evoked DA bursts in selective striatal subdivisions (Stelly, Tritley, Rafati, & Wanat, 2020). Dopamine’s enhancement via L-DOPA administration, moreover, improves reward but not action cost learning (Skvortsova et al., 2017). To the degree that pupillometry can be considered a proxy measure of activity of ascending neuromodulatory systems, these findings are consilient with greater encoding of RPEs by pupil size fluctuations during acute stress. Negative correlations between SBP and PANAS negative affect and RPE-pupil size slopes suggest that primarily moderately stressed participants displayed a preference for maximizing reward value, which might be consistent with an inverted U-shape relationship between cognitive performance and DA transmission which is modulated by stress (Arnsten & Goldman-Rakic, 1998; Baik, 2020). Noradrenaline, however, mobilizes available energy to complete effortful actions and *locus coeruleus* neurons track energy expenditure (Varazzani et al., 2015). Stress-induced sAA concentrations, increased heart rate, and group differences in the association between pupil size fluctuations and EPEs all point to the involvement of the noradrenaline system. Thus, our model-based pupillometry and stress-induction results hint at stress-sensitive dopaminergic and noradrenergic mechanisms that may regulate cost and benefit learning, which could be explored in future work using targeted pharmacological approaches.

The results presented here may improve understanding of stress-related psychopathology. While asymmetric cost-benefit learning during acute stress may be beneficial to reach a desired goal state (e.g., safety) despite high action cost, such strategies could also be maladaptive. For example, stress exposure can lead to drug or smoking relapse (L. Schwabe, Dickinson, & Wolf, 2011), a context in which reward value and action cost may be misaligned. Cost-benefit reinforcement learning may provide a useful framework to test hypotheses regarding stress-related impairments in learning and decision-making.

Some study limitations need to be acknowledged. First, pupil dilation associated with effort expenditure greatly reduced our power to detect robust associations between EPE encoding and pupil size fluctuations. Future studies should, therefore, consider a temporal delay between effort outcome and effort expenditure phases. Second, while our computational model was able to recover overall task performance patterns in both groups, such effects were subtle and dependent on the contribution of other (non-learning) parameters, which may highlight the importance of interindividual differences in model parameters.

To summarize, we present evidence of asymmetric effects of acute stress on cost versus benefit reinforcement learning during acute stress, which computational analyses of task behaviour explain as a failure to assign appropriate importance to RPEs versus EPEs, and our model-based pupillometry tentatively link to activity of ascending midbrain neuromodulatory systems. These results highlight for the first time how learning under acute stress can be tilted in favour of acquiring good things and away from the avoidance of costly things.

## Materials and Methods

### Participants

Adult participants were recruited via paper and online advertisements. All participants were screened for a DSM-5 psychiatric and/or neurological disorder, substance use, endocrine and/or vascular disorder, abnormal BMI (>40 or <18), smoking and drinking (>10 cigarettes/units per week), psychotropic medication use (lifetime) and hormonal contraceptive use (current; female participants only). All participants completed the ∼2-hour experiment between 12:00h and 18:00h to minimize diurnal cortisol fluctuations (Bailey & Heitkemper, 2001). Participants were instructed to refrain from alcohol (starting the evening before the day of the experiment), smoking, food, caffeine intake, strenuous physical activity and brushing their teeth (all >2 hr prior to experiment), which was verified verbally at the start of the session. Four participants were excluded due to an equipment failure (n=4). Three participants quit during stress-induction (n=2) or task procedures (n=1). Because chance-level performance on reinforcement learning tasks might indicate a successful manipulation, a lack of motivation, or a failure to comprehend the task instructions, participants that performed at or below chance level (0.5) on both RL and/or both EL pairs near the end of the experiment (final 10 presentations) were excluded (n=13; 6 acute stress, 7 no-stress control). Including these participants did not alter our key finding that acute stress was associated with asymmetric cost versus benefit learning (see Results). Pupillometry and neuroendocrine data were not processed further for these participants. The study was approved by the ethics committee of the Faculty of Psychology and Neuroscience, Maastricht University (ERCPN-197_03_08_2018) and carried out in accordance with the Declaration of Helsinki. Participants were remunerated in gift vouchers or research participation credits. Task earnings were paid out in gift vouchers.

### Acute stress induction

The MAST is a validated stress-induction paradigm combining both psychological and physiological stressors, and robustly increases neuroendocrine, physiological, and subjective indices of acute stress (Smeets et al., 2012). During a 5-min preparation phase, participants are instructed about the upcoming task via oral and visually displayed instructions, followed by a 10-min stress-induction phase consisting of alternating blocks of cold-water immersion (non-dominant hand; 2°C) and backward counting in steps of 17 (while receiving negative evaluative feedback from an experimenter), with a (non-recording) camera continuously directed at the participant’s face, which was displayed to the participant on a second display. During the MAST no-stress control condition, participants immerse their hand in lukewarm water (36°C) and perform simple mental arithmetic, e.g., counting from 1 to 25, without receiving feedback or fake camera recordings.

### Neuroendocrine, physiological and subjective stress measurements

sCORT and sAA were collected to measure stress-induced increases in hypothalamic-pituitary-adrenal (HPA) axis and sympathetic-adrenal-medullary (SAM) axis activity, respectively (Dickerson & Kemeny, 2004; Koh, Ng, & Naing, 2014). Saliva samples were obtained using synthetic Salivette® devices (Sarstedt, Etten-Leur, the Netherlands) during 3-min sampling periods at 6 time points. A baseline sample was collected 10 min prior to the MAST (baseline: t_1_ = t_-10_) and five samples post-MAST (t_2_= t_+00_, t_3_= t_+10_, t_4_= t_+20_, t_5_= t_+30,_ t_6_= t_+40_). SAA assessments were obtained only for t_1_-t_4_, due to the rapid decay of sAA post-stress induction (Dennis Hernaus, Quaedflieg, Offermann, Casales Santa, & van Amelsvoort, 2018; Nater et al., 2005). For all participants, t_2_ marked the starting point for the reward value maximisation/action cost minimisation task. Samples were stored at −20°C immediately after completion of each session. SCORT and sAA levels were determined using a commercially available luminescence immune assay kit (IBL, Hamburg, Germany) and kinetic reaction assay (Salimetrics, Penn State, PA, USA), respectively.

Systolic blood pressure (SBP) and heart rate (HR) as an index of autonomic nervous system (ANS) arousal (Schubert et al., 2009; Wright, O’Brien, Hazi, & Kent, 2014) were assessed at t_1_ and t_2_ using an OMRON M4-I blood pressure monitor (OMRON Healthcare Europe B.V., Hoofddorp, The Netherlands). Subjective affect ratings were assessed at t_1_ and t_2_ using the 20-item Positive and Negative Affect Scale (PANAS) (Watson, Clark, & Tellegen, 1988).

### Reward maximization versus action cost minimization reinforcement learning task

All participants completed a probabilistic stimulus selection paradigm during which they learned to select stimuli with high reward value (20 Eurocents) and avoid stimuli with high action cost (exerting force above a pre-calibrated individual threshold for a duration of 3000ms). This reinforcement learning task is conceptually similar to a previously-validated probabilistic action selection task that has been employed to study the neural signatures of reward and effort prediction errors and dopaminergic drug effects on reward-effort computations (Skvortsova et al., 2017; Skvortsova et al., 2014). The paradigm was designed in PsychoPy v3.0.0b11 (Peirce et al., 2019) and presented on a 24″ monitor (iiyama ProLite b2483HSU). Physical effort (in mV/kgf) was registered using a hand-held dynamometer in combination with a transducer amplifier (DA100C) and data acquisition system (MP160; all manufactured by BIOPAC Systems, Inc). Individual effort thresholds used throughout the task were obtained by calculating 50% of each participant’s maximal voluntary contraction (MVC) (Le Heron et al., 2018) reached over three calibration trials by squeezing the dynamometer with the dominant hand.

On each of 120 trials, participants chose between two paired distinct black-and-white images (“stimuli”) that were probabilistically associated with both the receipt of a monetary reward and exertion of physical effort (see Figure 1 for a graphical overview). At trial onset, a fixation cross flanked by two images was presented; participants chose one image by pressing the V/B button for the left/right option, respectively. A 440Hz/600Hz tone for left/right choice (200ms) was presented to confirm the participant’s choice. Next, a thermometer with the command “SQUEEZE” or “DON’T SQUEEZE” was displayed. If participants were required to exert effort, they were instructed to squeeze the dynamometer until the mercury level reached the top. The mercury bar only moved if participants exerted above-threshold levels of force and stopped moving if exerted force fell below. The cumulative above-threshold time was 3000ms. If no effort production was required, an animation of a rising mercury bar was displayed (3000ms). Finally, a screen was presented showing either a €0.20 coin or a crossed-out coin, indicating no reward (3000ms).

Participants learned to choose the optimal (most reward or most effort avoiding) stimulus for four distinct image pairs, 30 presentations each, with yoked reward and action cost contingencies. For 2/4 pairs, participants could regularly acquire rewards by selecting one (optimal) stimulus over the (suboptimal) other (henceforth, “reward learning”/RL pairs), while the probability of having to exert effort was identical for both stimuli. For the other two pairs, choices of one stimulus were more frequently followed by the avoidance of effort (“effort learning”/EL pairs), while the probability of reward was kept constant between both. For all pairs, the probability of the stimulus property that was *kept constant* (reward/effort) was set to a 33.3% chance of positive outcome upon selection (reward/ effort avoidance) and 66.6% chance of negative outcome (no reward/effort).

To assess whether any acute stress effects on reward maximization (measured using RL pairs) and effort cost minimization (measured using EL pairs) learning were potentially mediated by task difficulty, we employed different difficulty levels for each RL and EL pair. That is, for one RL and one EL (“easy”) pair, a choice for the optimal stimulus was followed by a positive outcome in 83% (vs. 17% negative outcome) of all trials (83% negative/17% positive outcome for suboptimal stimulus); for the other RL and EL (“hard”) pair a choice for the optimal stimulus was followed by a positive outcome in 70% (vs. 30% negative outcome) of all trials (and 70% negative/30% positive outcome) for the suboptimal stimulus. This approach allowed us to disentangle whether acute stress primarily impacted domain-specific (RL vs. EL) or general (easy vs. hard) reinforcement learning (the latter which also might involve other cognitive skills that might be beneficial to performance and sensitive to change under stress, such as working memory (Schoofs, Wolf, & Smeets, 2009). The task contingencies described above were based on extensive pilot tests to identify a reinforcement schedule that would enable us to detect stress-induced improvements *and* decreases in task performance. We selected task contingencies based on pilot sessions involving a no-stress control condition and chose a reinforcement schedule associated with non-ceiling/floor performance on RL and EL trials.

Following the learning phase, participants completed a surprise test phase, similar to previous work (D. Hernaus et al., 2019; D. Hernaus et al., 2018). This phase consisted of 64 trials in which participants were presented with the original four, as well as six novel, stimulus combinations. Participants were asked to choose the stimulus with the highest reward value or the lowest action cost - depending on a coin or thermometer image presented in the middle of the screen - and received no choice feedback. This allowed us to assess acquired choice tendencies, as well as generalizability of this information to novel situations. The four original pairs were presented four times (total n=16) during which we only asked participants to discriminate on the basis on the reward value (for RL) or action cost (for EL). For novel stimulus combinations, we only presented stimuli that differed in reward value/action cost if reward value discrimination/action cost discrimination was assessed (total n=48: n=4 presentations for the 6 combinations).

For every participant, stimuli were randomly assigned to pairs, optimal/suboptimal stimulus orientation was balanced (50% of all optimal stimulus presentations occurred on the left-hand side) and misleading outcomes (e.g., negative outcomes for optimal stimuli) were equally spaced out across the thirty presentations (and balanced for left/right side). Trial presentation order was pseudo-randomized such that I) a given pair would never be presented more than twice in a row and II) the gap between two presentations of a given pair was never greater than four trials.

Prior to performing the actual task and prior before acute stress/no-stress control procedures, participants received standard verbal instructions and completed a 16-trial practice round of the learning phase. Participants were not informed about stimulus-outcome contingencies; they were only advised to accrue as much money as possible and avoid exerting unnecessary effort. A 60% accuracy performance threshold was used to confirm that participants understood the general task procedure. The practice round was repeated if participants failed to reach 60% accuracy. To prevent learning, we used deterministic stimulus-outcome probabilities and different stimuli.

### Computational cost-benefit reinforcement learning model: model space

In an attempt to uncover latent mechanisms by which acute stress affects reward maximization and/or action cost minimization, we turned to cognitive computational modelling. We employed a modified reinforcement learning framework based on Rescorla and Wagner (Rescorla, 1972), and used in Skvortsova et al. (Skvortsova et al., 2017; Skvortsova et al., 2014) to investigate whether acute stress impacted learning about sensitivity to, and/or discounting of reward value and action cost. We first describe the model space.

Various reinforcement learning models assume that choice preferences of an agent are updated via the prediction error, i.e., the mismatch between outcome and expectation (equation 1A, 1B) and the critical quantity that drives learning (Rescorla, 1972):

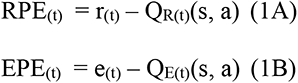

Here, Q_R(t)_(s, a) and Q_E(t)_(s, a) represent the expected reward value and action cost (i.e., effort), where *s* reflects the given pair and *a* refers to the more abstract action of selecting a stimulus (not to be confused with action selection), r_(t)_ and e_(t)_ represent the reward and effort outcome for the chosen stimulus at trial *t*. RPE_(t)_ and EPE_(t),_ thus, represent the RPE and EPE at trial t, respectively.

In order to allow for the possibility that humans do not calculate the prediction error against the actual outcome but, rather, what the outcome “feels” like (Huys et al., 2013), we considered a scenario in which reward and effort outcomes are first multiplied by a free parameter that captures the weight that reward and effort outcomes receive (“W_R_” and W_E_” in equation 2A and 2B). As the value of these parameters approaches 1, rewards are increasingly valued more positively, and effort more negatively. These parameters, therefore, control the maximum size of the prediction error.

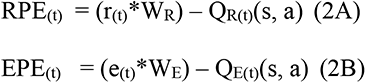

In various formulations of reinforcement learning, such as Q-learning (Watkins & Dayan, 1992) and the actor-critic framework (Niv, 2009; Rescorla, 1972), the degree to which prediction errors update choice preferences is represented by α, the learning rate (equation 3A), which determines how current prediction errors update choice preferences on the subsequent trial. High values of *α* allow for rapid updating of choice preferences, while a low *α* implies that choice preferences are updated at a slower pace and are thus co-determined by outcomes further into the past.

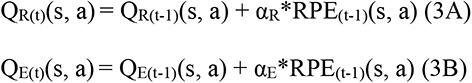

Extensive evidence suggests that organisms use different learning systems for different types of information, including reward value and action cost (Palminteri & Pessiglione, 2017; Skvortsova et al., 2017; Skvortsova et al., 2014) (equation 3A/B). Thus, the use of separate learning rates for RPEs and EPEs allows for asymmetrical learning about these types of information.

While the learning rate controls the *speed* at which choice preferences are updated, learning rate (nor reward/effort weight) alone does not explain how learned estimates of reward value and action cost may compete at the decision stage (i.e., when participants choose between two stimuli). Agents weight costs against benefits to calculate a subjective decision value (Pessiglione et al., 2017; Skvortsova et al., 2017), which is used to guide choices (equation 4).

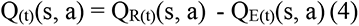

In its simplest form, Q, the subjective decision value of a stimulus is represented by the difference between the expected reward and action cost value at trial *t* (equation 4) (Skvortsova et al., 2014). However, this particular operationalization of subjective value does not take into account the observation that humans tend to discount or prioritize certain types of information in their decisions (Apps, Grima, Manohar, & Husain, 2015; Inzlicht et al., 2018). We, therefore, allowed for variation in the calculation of subjective decision value via action cost discounting (equation 5). While discounting rates can be linear or hyperbolic (Hartmann, Hager, Tobler, & Kaiser, 2013), here we only considered linear discounting in light of previous work using a similar task design (Skvortsova et al., 2017; Skvortsova et al., 2014). As the value of γ approaches zero, action cost discounted increases leading the agent to ignore action cost/only utilize reward value to make a decision.

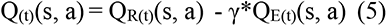

Once the subjective decision value has been computed, the degree to which participants deterministically sample the optimal stimulus is captured by a softmax decision function (equation 6).

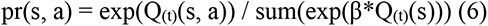

Here, pr is the probability of selecting an action, β is the inverse temperature parameter that among others captures the balance between exploration and exploitation (Nassar & Frank, 2016), Q_(t)_(s, a) is the net value of the chosen option and Q_(t)_(s) represents the net values of both stimuli in the pair.

Within the above-described model space our predictions of acute stress effects on reward maximization and action cost minimization could, thus, be explained by changes in sensitivity to reward value and/or action cost (W_R,_ W_E_), changes in how much weight RPEs and EPEs are afforded (i.e., learning rates, α_R,_ α_E_), and/or changes in the discounting of reward value by action cost (γ). If acute stress leads to more random responses, such effects should be captured by β.

Based on our predictions and the obtained pattern of results (most notably asymmetrical RL/EL performance in the acute stress condition), we considered six candidate models that could capture these various scenarios: I) a model with 2 distinct learning rates for reward and effort (αR, αE) [2LR]; II) a model with 2 learning rates (α_R_, α_E_) and a discounting parameter (γ) (2LR_ γ); III) a model with 2 learning rates (α_R_, α_E_), a reward weight (W_R_) and an effort weight parameter (W_E_) (2LR_W_R__W_E_), IV) a model with a *single* learning rate (α), reward weight (W_R_), effort weight (W_E_), and a discounting (γ) parameter (LR_ W_R___W_E__ γ); V) a model with 2 learning rates (α_R_, α_E_), a reward weight (W_R_), and a discounting (γ) parameter (2LR_W_R__γ); VI) a model with 2 learning rates (α_R_, α_E_), a reward weight (W_R_), effort weight (W_E_) and discounting (γ) parameter (2LR_ W_R__ W_E__ γ).

Lower/upper bounds for all parameters were set to [0,1] and all models contained a β parameter. Consistent with previous work (Skvortsova et al., 2017; Skvortsova et al., 2014), reward and action cost outcomes were set to [0,1 for no/yes reward] and [-1,0 for no/yes effort avoidance], respectively.

### Pupillometry

Fluctuations in pupil diameter were continuously measured using an SR-Research Eyelink 1000 Tower Mount infrared eye tracker while participants performed the reward maximization/action cost minimization reinforcement learning task (1000Hz sampling rate, except for three participants, whose data were obtained at 500Hz). Participants placed their head on an adjustable chin rest and against a forehead bar to minimize motion. Eye-tracker calibration was performed at the start of the paradigm, and subsequently every 10 min. Stimulus luminance was matched using the SHINE toolbox (Willenbockel et al., 2010) in MATLAB (v. 2014B; The MathWorks, Inc., Natick, Massachusetts, United States). Due to the COVID-19 pandemic, pupillometry data were not collected for the final eight participants. Three participants, moreover, failed the quality control for eye-tracking data (2 no-stress control/1 acute stress) leaving a final sample of 69 participants with eye-tracking data (34 no-stress control/35 acute stress).

Eye-tracking data were pre-processed using an open source pre-processing toolbox (Kret & Sjak-Shie, 2019) and in accordance with previous work (Jackson & Sirois, 2009). Blinks and other invalid samples, due to dilation speed, deviation from the trend line, and extreme values (Kret & Sjak-Shie, 2019) were removed, interpolated, smoothed (4Hz low-pass filter, fourth-order Butterworth filter) (Jackson & Sirois, 2009), z-scored and down-sampled to 50hz (i.e., 20ms). Bins with fewer than 80% valid samples were removed (Lawson et al., 2020). For analyses, we considered three epochs of interest: choice (−1500ms pre-choice - 15000ms post-choice), effort outcome (0-1000ms post-outcome), and reward outcome (0-2000ms post-outcome). We reduced the duration of the effort outcome epoch to 1000ms to minimize force exertion-related effects on pupil size (see below). Recent work has shown that expectation violations (prediction errors) are encoded by pupil size fluctuations within this timeframe (Lawson et al., 2020). Given that we observed large grip force-associated effects on the pupillometry signal (see Figure Supplement 8 middle row, for a comparison between effort and effort avoidance trials), we limited effort outcome analyses in the main text to effort avoidance trials, although we also report analyses involving all effort outcome trials in Figure Supplement 8.

### Statistical analyses

Statistical analyses were conducted using R, version 3.6.2 (Team, 2020) and, where applicable, results were visualised using Raincloud Plots (Allen, Poggiali, Whitaker, Marshall, & Kievit, 2019). Acute stress measurements were analysed using mixed ANOVAs involving Condition (between-factor condition: no-stress control, acute stress induction) and Time (within-factor: 2 pre/post-MAST or 6 levels for sCORT).

For the reward maximisation/action cost minimisation reinforcement learning task, an accuracy score was calculated by dividing the number of optimal stimulus choices by the total trial amount (*n*=30 per pair). Mixed ANOVAs involving Condition, Trial Type (RL, EL) and Difficulty (Easy, Hard pairs) were carried out. For analyses involving Time effects (i.e., repeated presentations of stimulus pairs), accuracy scores were averaged per bin of ten presentations (presentation 1-10, 11-20, and 21-30). To better understand whether acute stress effects on task performance were primarily driven by changes in sensitivity to positive or negative outcomes, win-stay (repeating a choice following a positive outcome) and lose-shift (choosing the other stimulus following a loss) rates were calculated for RL and EL trials (Hanneke E. M. den Ouden et al., 2013). For RL trials, we calculated win-stay/lose-shift rates using reward outcomes (yes/no reward); for EL trials we used effort outcomes (yes/no effort). We refer to the 2-level factor representing win-stay/lose-shift rates as “Strategy”. For surprise test trials involving the original four pairs (n=4 presentations per pair), we investigated final choice tendences using a one-sample t-test against chance level (0.5). Participants’ ability to discriminate stimuli based on reward value and action cost in novel stimulus arrangements (n=48, 24 reward value and 24 action cost discrimination trials) were investigated using mixed ANOVAs involving Condition and Trial Type.

Group differences in model parameters from the non-hierarchically fit model were investigated using Condition-by learning rate (α_R_, α_Ε_) mixed ANOVAs and independent samples t-tests. Given that we used separate priors for the two groups, we report the Bayesian analogue of a t-test and mixed-ANOVA (Kruschke, 2014) - a more robust test of group differences - for posterior parameters obtained from the hierarchically fit model (for reference, we also report these analyses for the non-hierarchical data).

*Post hoc* (simple) main effect analyses for all ANOVAs were conducted using independent sample (Condition), paired-samples (Time, Trial Type, Strategy), and one-sample t-tests (≠ 0 or 0.5). Greenhouse–Geisser-corrected statistics were reported when sphericity assumptions were violated. We report statistical significance as *p*<0.05 (two-sided), but we note that most main and interaction effects involving Condition survived at a more stringent threshold (*p*<.01), except for some strategy and surprise test phase effects, which should be interpreted with caution. In case of statistically significant results, generalized eta square (ges; n^2^_G_) was reported, with n^2^_G_ values of 0.02, 0.06, and 0.14 representing a small, medium, and large effect size, respectively (Lakens, 2013).

With respect to pupillometry, we conducted model-free and model-based analyses. In model-free analyses, we investigated group differences in pupil size during the choice, effort outcome, and reward outcome stage, for every bin of interest. To better understand how putative activity of ascending neuromodulatory systems may drive stress-induced changes in computational strategies that support reward maximisation/action cost minimisation learning, we conducted model-based pupillometry analyses using computational parameters from the winning (2LR_ γ) model (Lawson et al., 2020). First, we used linear regression to estimate beta weights for the association between pupil size and computational estimates of task-related behaviour for every participant, for every epoch, for every bin. For the choice phase, we regressed trial-wise measures of pupil size against trial-wise estimates of the subjective decision value (i.e., effort-discounted reward value) of the chosen stimulus. For the effort and reward outcome phase, trial-wise EPEs and RPEs were the primary predictors of interest, respectively. Trial number (1-120) and presented images/pair (RL_easy, RL_hard, EL_easy, EL hard) served as additional predictors of interest for all models. Additional epoch-specific variables of interest were included for the choice (optimal choice yes/no), effort (action cost of chosen stimulus, effort avoidance yes/no), and reward (reward value of chosen stimulus, reward yes/no, effort avoidance yes/no) outcome phase. Similar results were obtained when repeating the analyses with more elaborate GLMs (e.g., the addition of yes/no most likely outcome based on reward/effort outcome probabilities [“surprise”] and reward/action cost for EL/RL trials). Secondly, in group-level GLMs, we compared the resulting beta weights I) against zero (for the no-stress control/acute stress condition separately), to investigate when the pupil encoded the computational process of interest, and II) between groups, to assess stress-induced changes in associations between pupil size and computational processes. To control the false positive rate, we conducted permutation tests at the bin- and cluster-level (2000 permutations, *α*_permute_=0.05). All correlations were performed using Spearman’s ρ correlations. Permutation tests were also conducted for correlation analyses involving acute stress measures and pupil encoding of predictions errors.

**Figure 3 – Source Data 1**

**Source files for task performance data.**

This link contains all task performance data used for the analyses shown in Figure 3. Raw data can be found under “task_performance”.

https://osf.io/ydv2q/

**Figure 6 – Source Data 2**

**Source files for pupillometry data.**

This link contains pupillometry data used for the analyses shown in Figure 6. Raw data can be found under “pupillometry”.

https://osf.io/ydv2q/

## Acknowledgements

We are indebted to Drs. Conny Quaedflieg, Edwin S. Dalmaijer. and Ross D. Markello for advice on stress-induction procedures and paradigm development. We thank Dr. Michael Frank for advice on hierarchical computational modelling procedures. We thank Katya Brat-Matchett for involvement in participant recruitment and Truda Driessen for administrative support.

## Supplementary Figures

**Figure Supplement 1.**
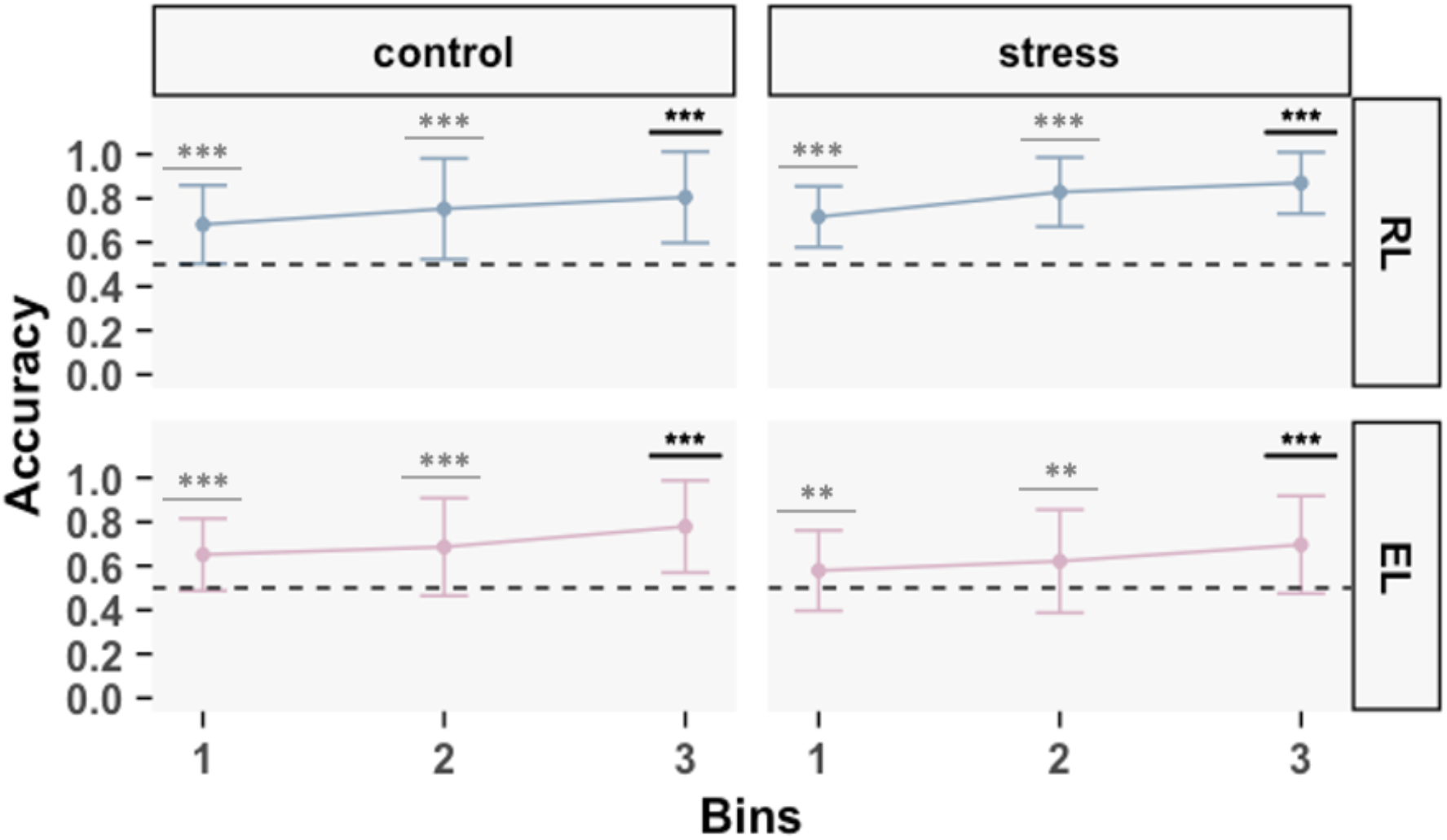
Evidence of reward and action cost reinforcement learning. Optimal stimulus choices (“accuracy”) on reward learning (RL) and effort learning (EL) (rows) trials for both conditions (columns). Trials were binned into groups of 10 presentations. Participants performed significantly better than chance level in all bins. Means ± SD. Significant differences are denoted by asterisks (*: *p* < 0.05, **: *p* < 0.01, ***: *p* < 0.001).

**Figure Supplement 2.**
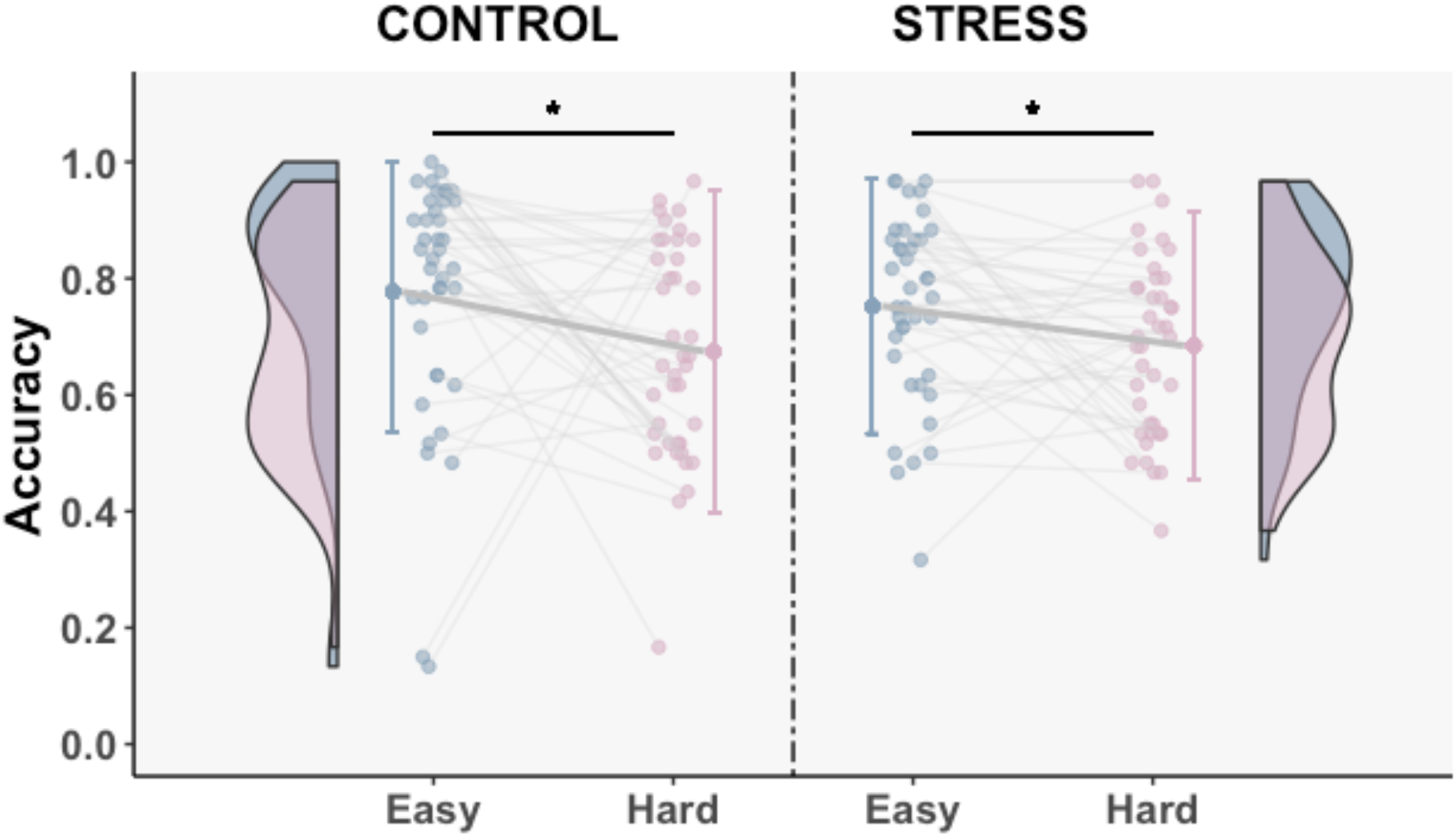
Acute stress does not affect difficulty learning. Easy and hard pairs collapsed across RL/EL trials depicted for each condition separately. While all participants sampled the optimal choices more frequently for Easy vs. Hard pairs, no significant Condition-by-Difficulty interaction or between-group differences were observed. Means ± SD, individual data points, distribution and density of the data are displayed. Significant differences are denoted by asterisks (*: *p* < 0.05, **: *p* < 0.01, ***: *p* < 0.001).

**Figure Supplement 3.**
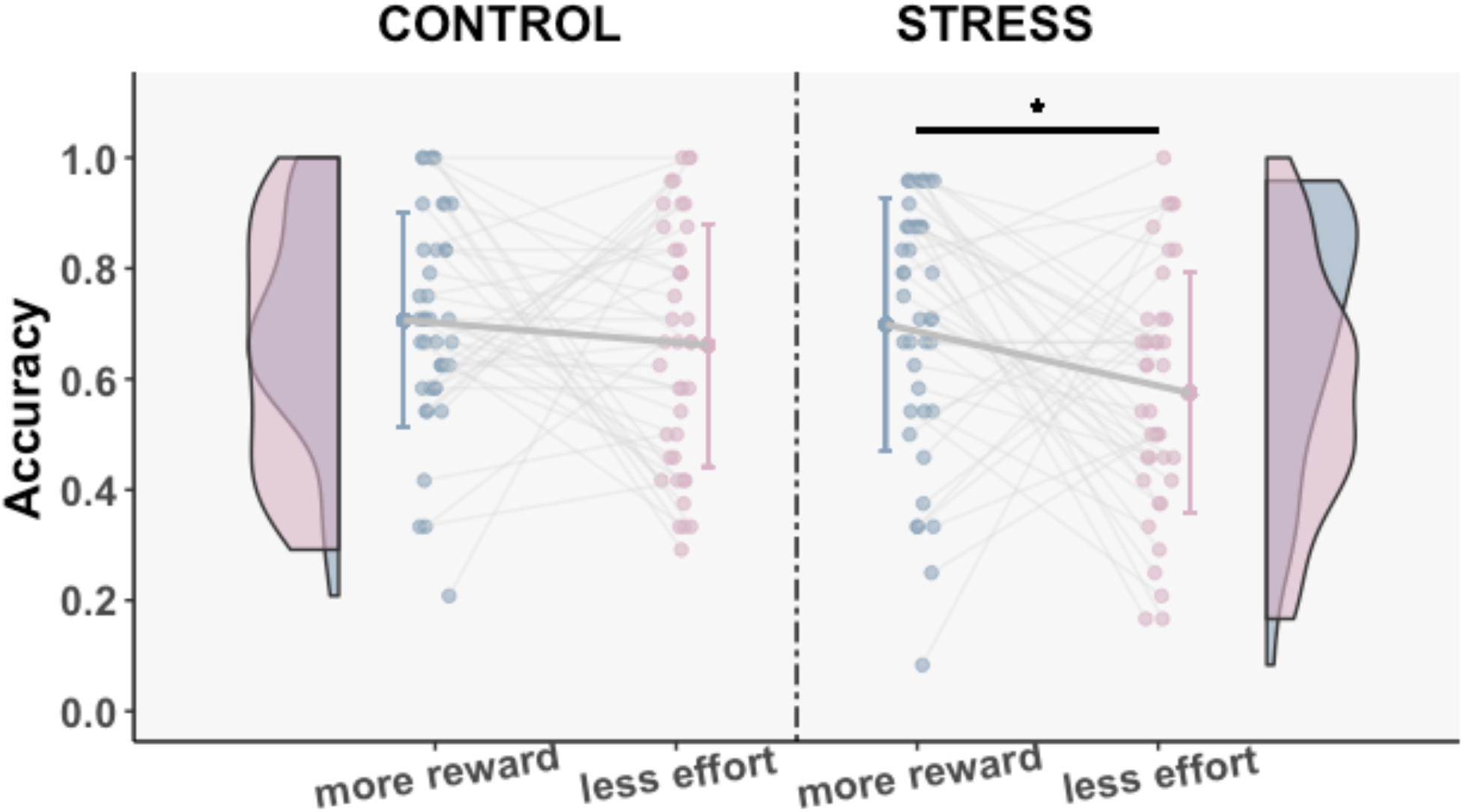
Surprise test phase performance. The acute stress group performed better on reward than action cost discrimination trials. Means ± SD, individual data points, distribution and density of the data are displayed. Significant differences are depicted with asterisks (*: *p* < 0.05, **: *p* < 0.01, ***: *p* < 0.001).

**Figure Supplement 4.**
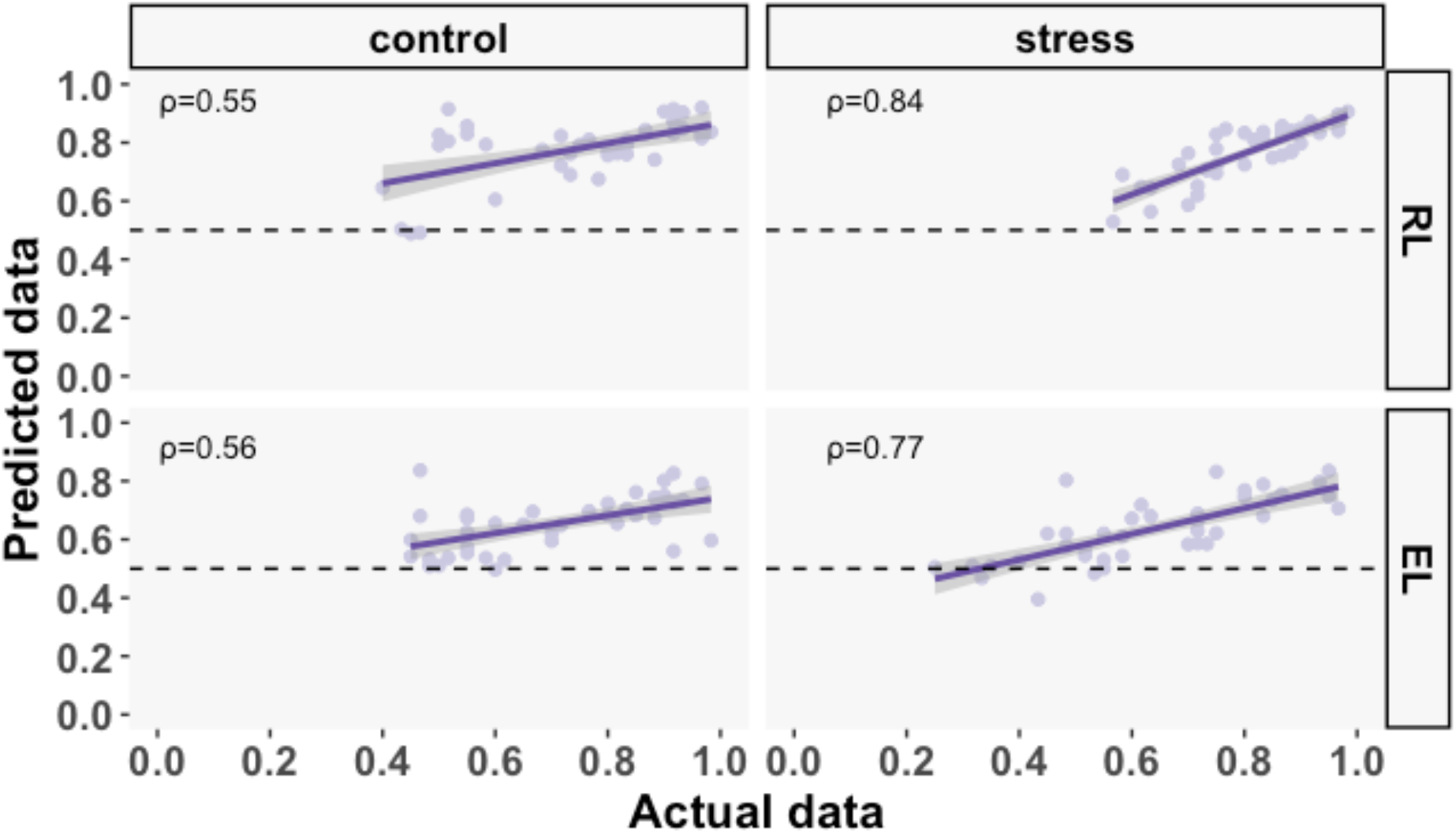
Correlations between empirical and simulated 2LR_γ choices. Actual and *post hoc* simulated choices for RL and EL (rows) were highly correlated both for no-stress control and acute stress subjects (columns). Simulations were averaged across 10 repetitions per subject. Solid and shaded lines represent mean ± CI_95%_. Dots represent individual data points. Horizontal dashed lines indicate chance level (0.5).

**Figure Supplement 5.**
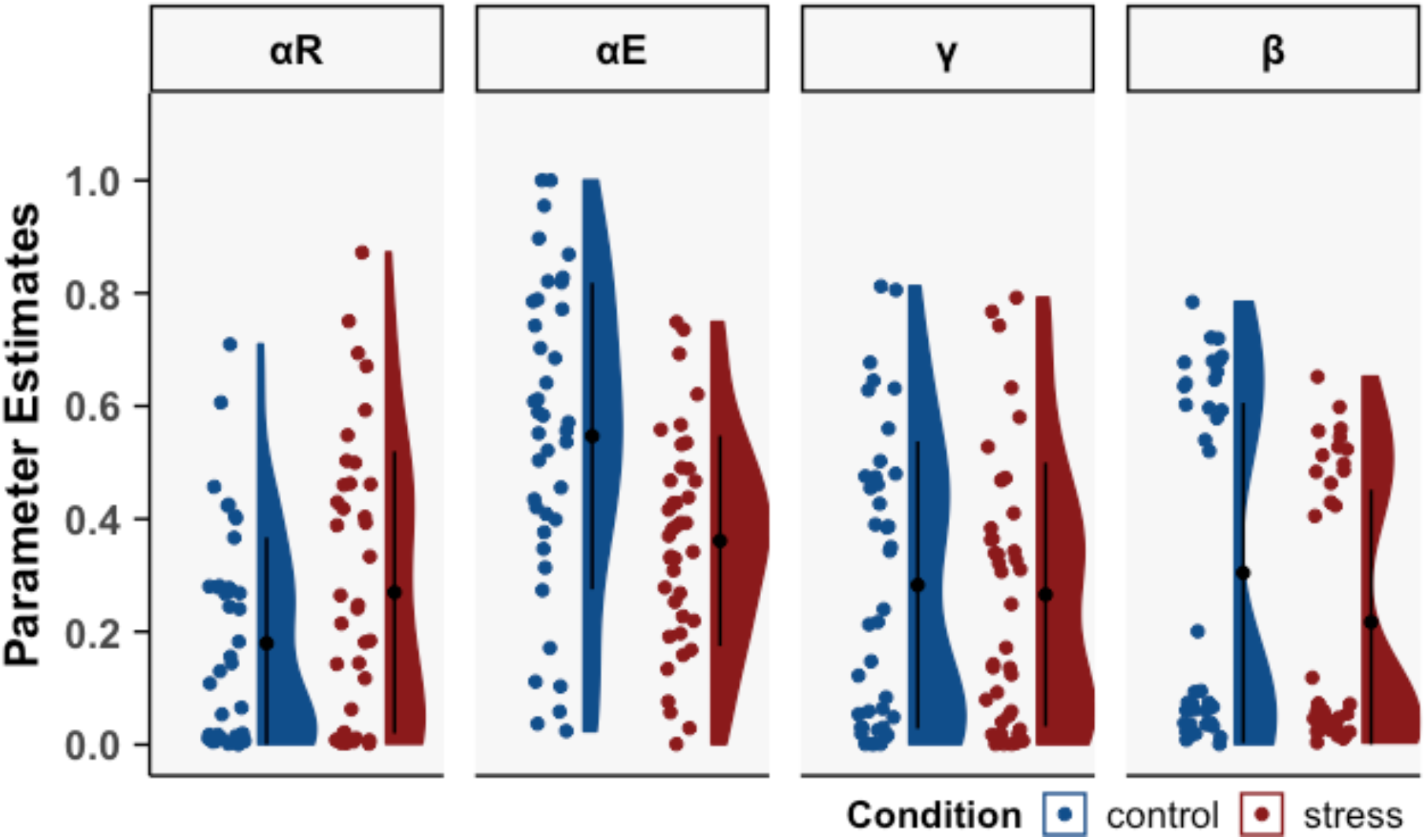
Parameter estimates after Bayesian hierarchical model fitting. Hierarchical model fitting reproduced the overall pattern of parameter estimates (Figure 5 for comparison).

**Figure Supplement 6.**
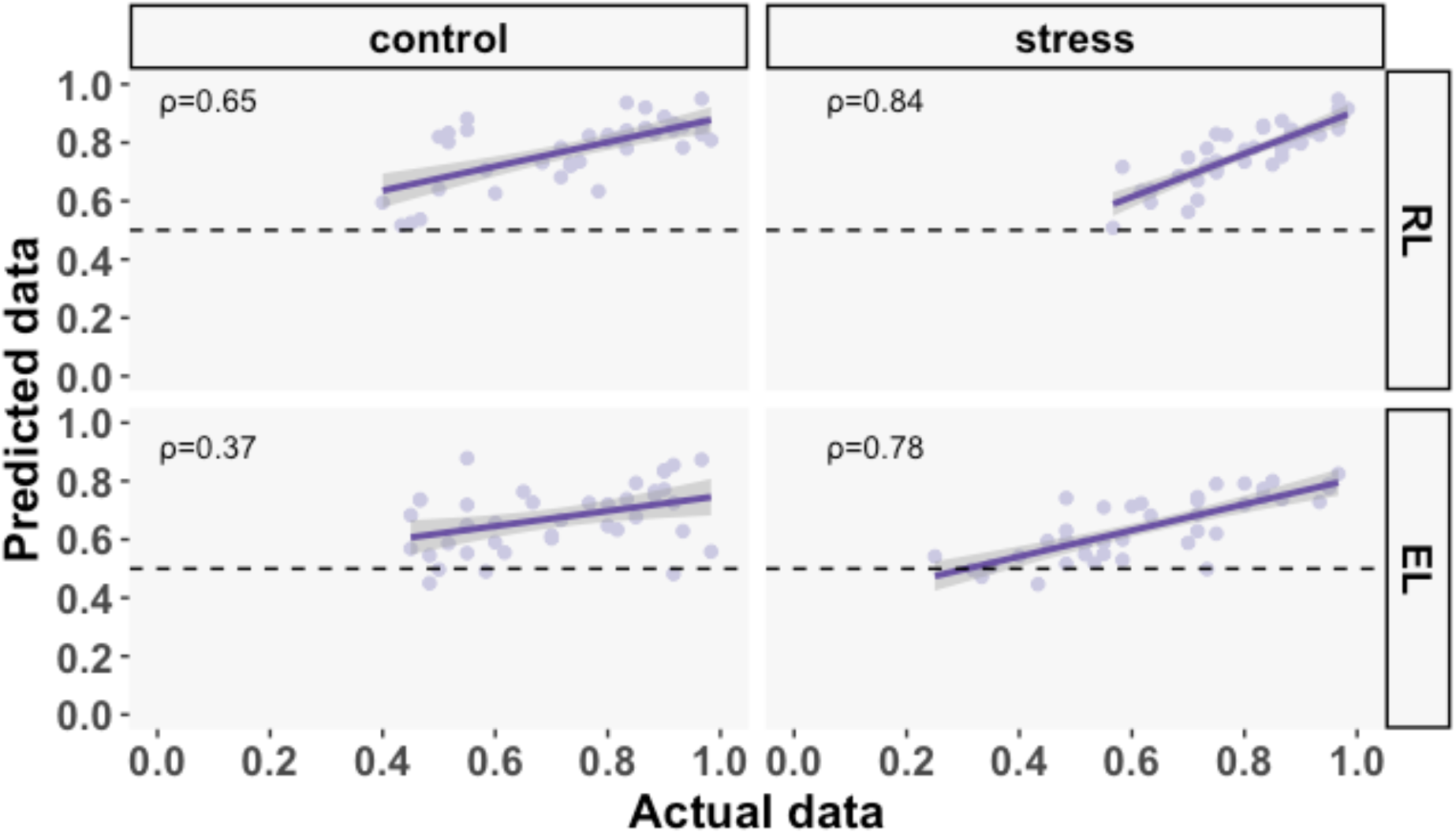
Correlations between empirical and simulated 2LR_γ choices after Bayesian hierarchical model fitting. Correlations between actual and *post hoc* simulated choices for RL and EL (rows) for no-stress control and acute stress subjects (columns). Simulations were averaged across 10 repetitions per subject. Solid and shaded lines represent mean ± CI_95%_. Dots represent individual data points. Horizontal dashed lines indicate chance level (0.5).

**Figure Supplement 7.**
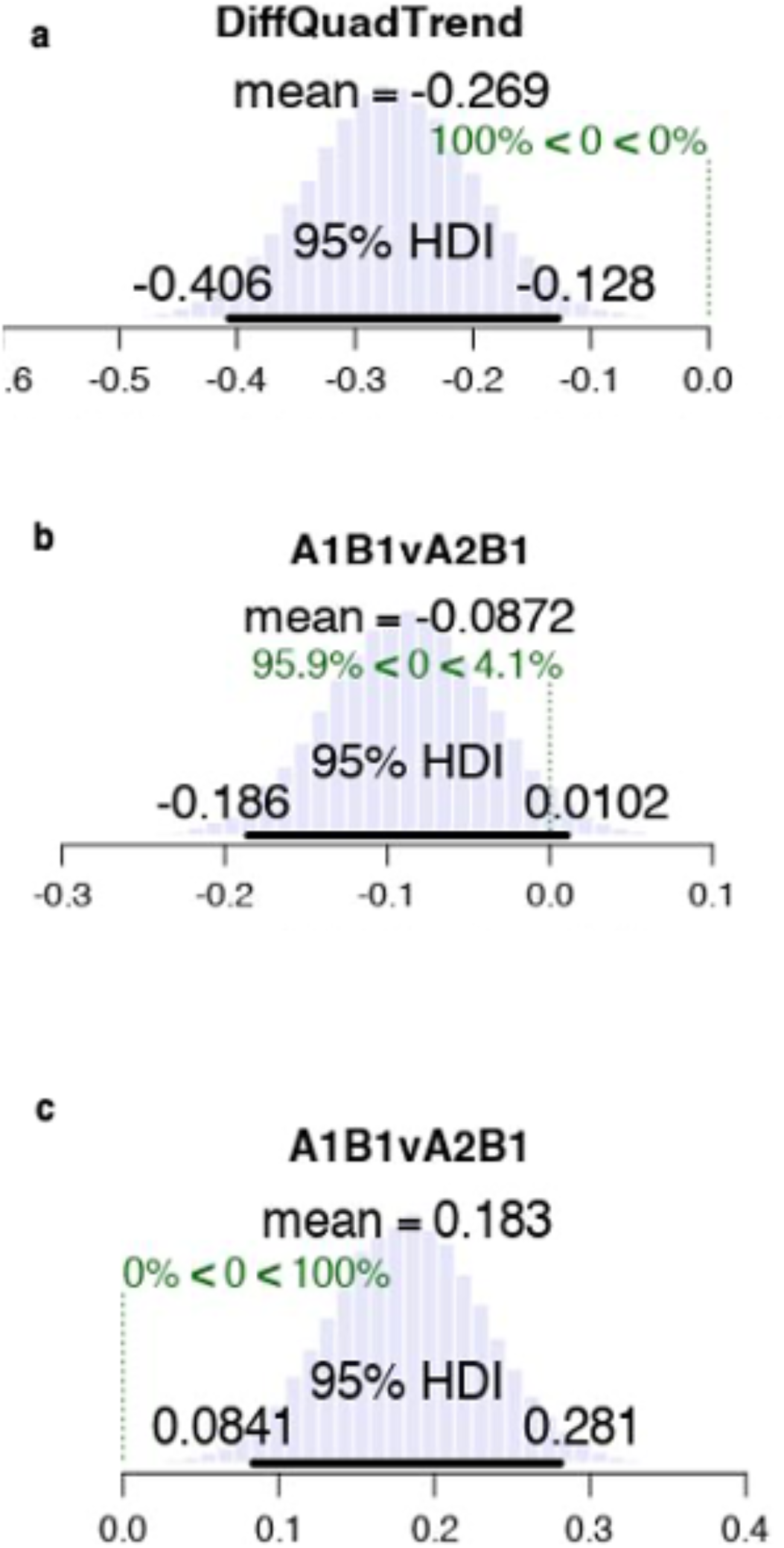
Bayesian estimation analysis to evaluate group differences in posterior parameter distributions. Panel A. Bayesian estimation (mixed-ANOVA) using posterior parameters (following hierarchical fitting) revealed evidence for a credible Condition-by-Learning Rate interaction. The observed mean difference from zero that falls outside the 95% HDI suggests that the difference between α_E_ and α_R_ was greater in no-stress controls compared to acute stress subjects. Panel B. Both groups did *not* differ in the magnitude of α_R_, as indicated by a 95% HDI that included 0. Panel C. Acute stress compared to no-stress control subjects exhibited a lower value of α_E_, as indicated by a 95% HDI that falls well above zero.

**Figure Supplement 8.**
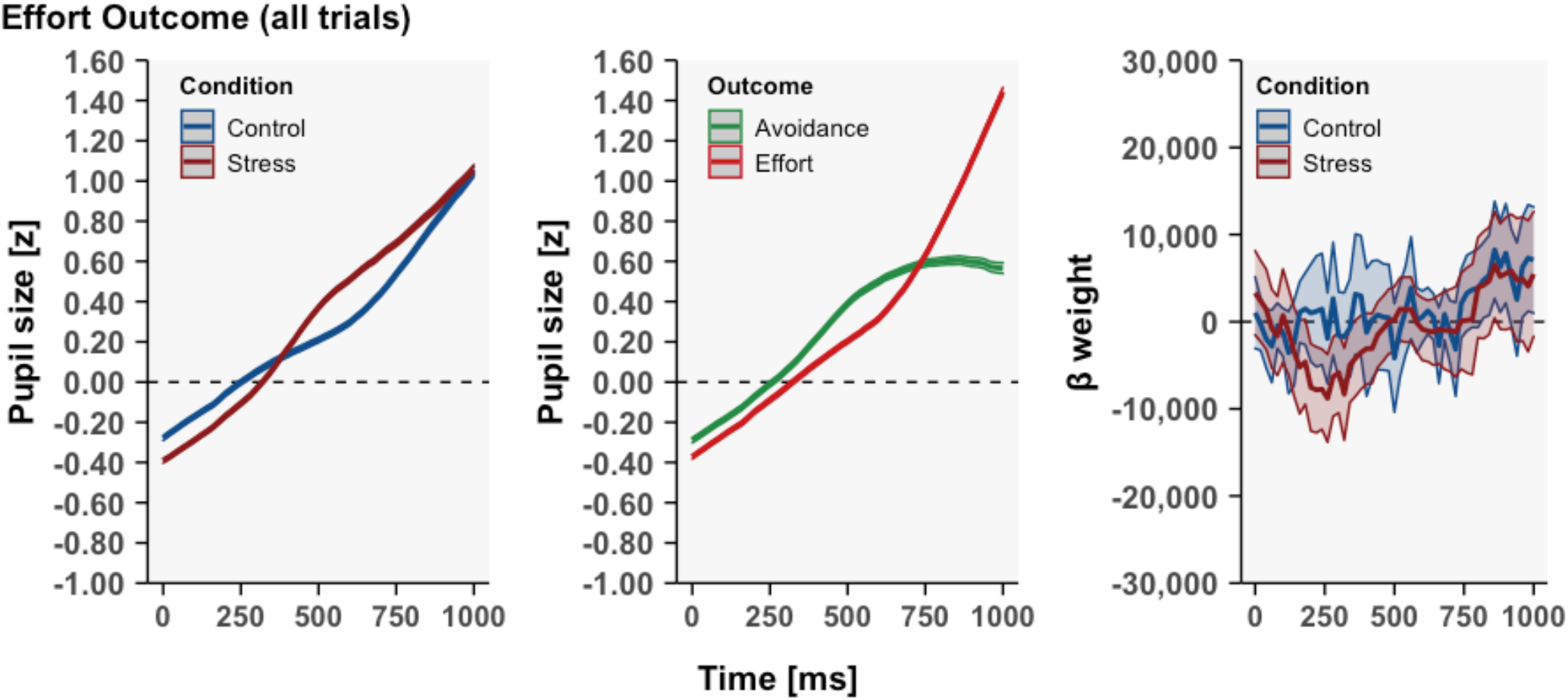
Pupillometry analyses using all effort outcome trials. **Left:** Model-free analyses of pupil size using all effort outcome trials. **Middle**: Pupil size differences during effort/effort avoidance outcomes in the entire sample; force exertion was associated with large effects on pupil size and these trials were therefore excluded from analysis. **Right**: Model-based action cost prediction error analyses using all effort outcome trials.

